# “Design principles of a membrane-spanning ubiquitin ligase”

**DOI:** 10.1101/2025.09.11.675358

**Authors:** Carys Williams, Laura M. Nocka, George Hedger, Pragya Parashara, Els Pardon, Naomi R. Latorraca, Ganesh V. Pusapati, Dorothy Lartey, Lei Gao, Ljiljana Milenkovic, Rod Chalk, Jan Steyaert, Susan Marqusee, Loïc Carrique, J. Fernando Bazan, Sarah L. Rouse, Jennifer H. Kong, Christian Siebold, Rajat Rohatgi

## Abstract

Receptor-type E3 ubiquitin ligases are membrane-spanning assemblies that enable extracellular signals to directly control ubiquitylation in the cytoplasm. Despite playing widespread roles in tissue patterning and homeostasis, metabolism, and immunity, their structures and mechanisms remain poorly understood. Using cryo-electron microscopy, integrated with biophysical and functional studies, we visualized an E3 complex composed of two transmembrane proteins, MEGF8 and MOSMO, and the intracellular RING-family protein MGRN1. This MEGF8-MOSMO-MGRN1 (MMM) complex regulates left-right patterning of the body axis and the development of multiple organs, partly by attenuating signaling through the Hedgehog pathway. We find that the MMM complex functions like a fishing pole: a long, flexible helix attached to a membrane platform suspends an activated and precisely oriented RING domain—like a fishhook—to ubiquitylate the cytoplasmic surfaces of target receptors. Our structure explains how mutations in *MEGF8* cause multi-organ birth defects in humans and defines a paradigm for receptor regulation by ubiquitylation.

## Introduction

The sensitivity of cells to extracellular ligands depends on the abundance of signaling receptors at the cell surface^1^. Increasing evidence suggests that receptor abundance at the plasma membrane can be controlled by membrane-spanning E3 ubiquitin ligases^2,3^. Like other E3s, these assemblies employ a RING (Really Interesting New Gene) domain to recruit and activate an E2-ubiquitin (Ub) conjugate, catalyzing Ub transfer from the E2 to the receptor substrate in the cytoplasm. Ubiquitylation leads to the removal of receptors from the plasma membrane and consequent loss of sensitivity to cognate ligands. However, this enigmatic class of E3s additionally contain transmembrane (TM) and extracellular domains, raising the question of how TM communication influences receptor recognition and ubiquitylation^2^.

Membrane-spanning E3s that target receptors are built around type I single-pass membrane proteins. The catalytic RING domain is either part of the same polypeptide chain as the TM helix or part of a separate protein that is recruited to the cytoplasmic tail of the type I membrane protein. Prominent examples of the former include ZNRF3 and RNF43, tumor suppressor proteins that constrain WNT, Bone Morphogenetic Protein (BMP) and Fibroblast Growth Factor (FGF) signaling by reducing the levels of their receptors at the cell surface^4–8^, and GRAIL/RNF128, an E3 that regulates receptors critical for T-cell responsiveness^9^. An example of a multi-subunit TM E3 is the heterotrimeric complex composed of MEGF8, a type I membrane protein with a large extracellular domain (ECD), MOSMO, a tetraspan membrane protein, and MGRN1, a cytosolic RING-type E3 ligase that stably associates with the intracellular domain (ICD) of MEGF8 (**Figures 1A and 1B**)^10–12^. This MMM (<u>M</u>EGF8-<u>M</u>OSMO-<u>M</u>GRN1) complex attenuates Hedgehog (HH) morphogen signal strength by ubiquitylating Smoothened (SMO), a G-protein coupled receptor (GPCR) that transduces the HH signal across the membrane. As a consequence, SMO is depleted from primary cilia, organelles where HH signaling is transmitted from the membrane to the cytoplasm^13,14^. Tight control of SMO activity is essential because HH signaling is involved in the development of nearly all human tissues. Excessive SMO activity causes birth defects and cancer^15,16^. Although SMO antagonists are approved for use in patients with cancers of the skin and blood, newer agents are needed due to resistance and side effects^17^. In addition, the MMM complex likely regulates signaling receptors other than SMO, since mutations in MMM genes lead to a wide range of severe birth defects in mice and humans^10,18^. MGRN1 also associates with Attractin (ATRN), a type I membrane protein related to MEGF8, to downregulate melanocortin receptors, GPCRs implicated in diverse physiological processes including pigmentation, food intake, and energy balance^19–23^. Thus, RING-type E3 ligases recruited to TM scaffolds play widespread (and largely unstudied) roles in regulating cell-to-cell communication.

**Figure 1.**
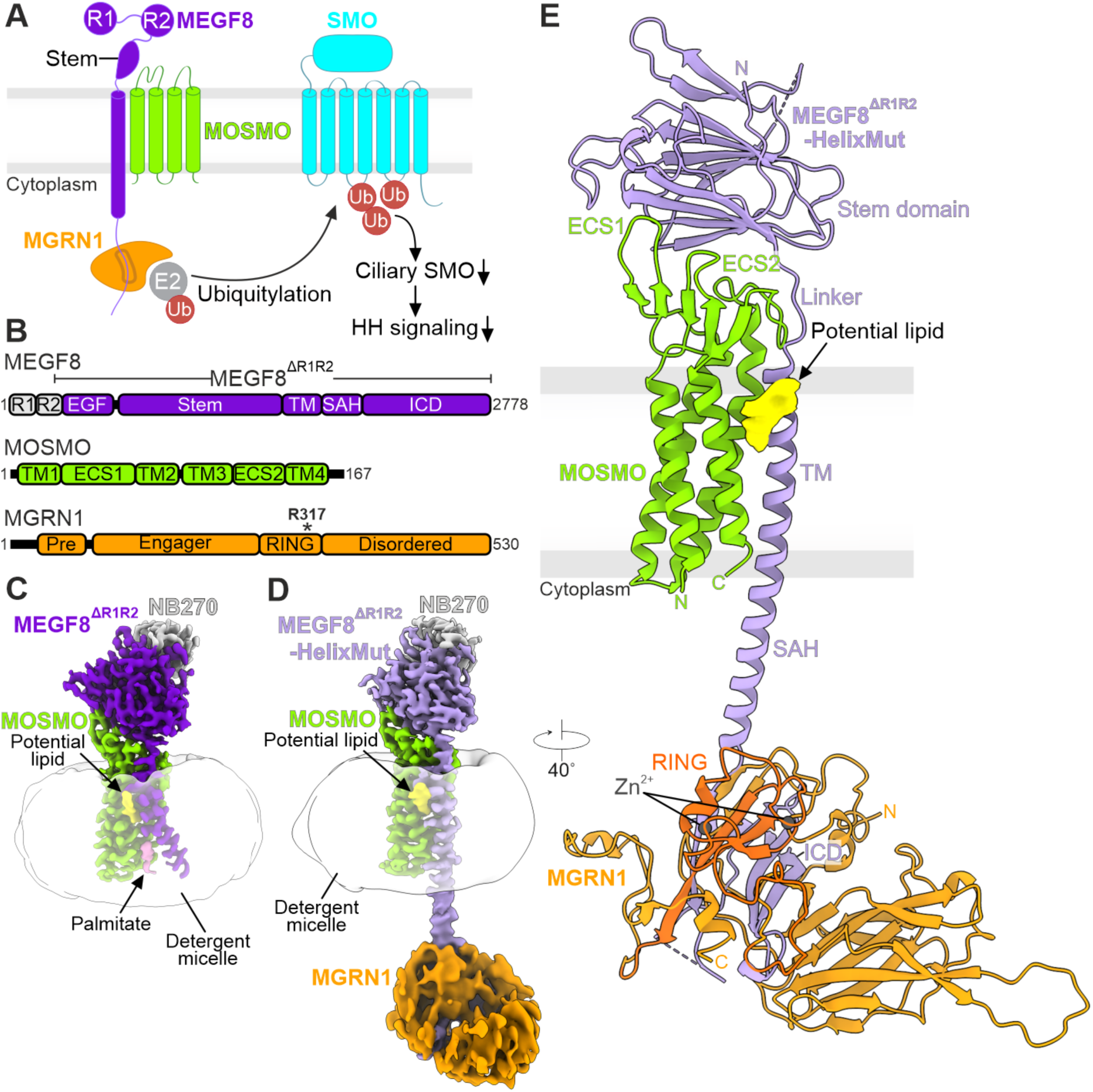
Architecture of the MEGF8-MOSMO-MGRN1 (MMM) complex. (**A** and **B**) Assembly (A) and domain composition (B) of the three components of the MMM complex, which attenuates HH signaling by ubiquitylating the GPCR and HH signal transducer SMO^10–12^. R1 and R2, large multi-domain pseudo repeats (Figure S1A); EGF, epidermal growth factor-like domain; TM, transmembrane helix; SAH, solitary α-helix; ICD, intracellular domain; ECS, extracellular segment; Pre, pre-engager domain. The asterisk shows the position of the linchpin residue in MGRN1 (R317) that contacts and activates the E2-Ub conjugate (B). (**C** and **D**) Cryo-EM maps of the MEGF8^ΔR1R2^-MOSMO-NB270 (2.65 Å, C) and MEGF8^ΔR1R2^-HelixMut-MOSMO-MGRN1-NB270 (average 3.3 Å, D) complexes shown in the same orientation. The SAH of MEGF8^ΔR1R2^ has been replaced by an ER/K helix (D, Figure S1A) to increase stability and to improve resolution of the intracellular components. (**E**) Cartoon representation of the helix-stabilized MMM complex. Color coding as in A, with the Zn^2+^-containing RING domain of MGRN1 highlighted in dark orange. The MEGF8 SAH suspends MGRN1 below the membrane and the MEGF8 ICD forms an intertwined structural unit with MGRN1. Cryo-EM map for a potential sterol is depicted in yellow. Termini are labeled. Zn^2+^ ions are shown as gray spheres. See also **Figures S1 and S2**.

Despite the essential roles of membrane-spanning E3s in development, tissue homeostasis, immune function, and metabolism, no structural information is available for these ubiquitin ligases in their membrane-embedded form– comprising extracellular, transmembrane, and cytoplasmic domains. The lack of three-dimensional structures has limited our understanding of how these multi-subunit complexes function at the membrane. To address this fundamental gap in knowledge, we investigated the architecture and molecular mechanism of the MMM complex as an exemplar of multi-subunit, membrane-spanning E3 ligases. Using cryo-electron microscopy (cryo-EM), we determined several high-resolution structures of the heterotrimeric MMM complex and characterized conformational dynamics through hydrogen-deuterium exchange mass spectrometry (HDX-MS) and atomistic simulations. Coupled with ubiquitylation and cellular signaling assays, our findings define a mechanistic blueprint for how TM scaffolds precisely suspend and orient RING domains beneath the inner leaflet of the plasma membrane to capture target receptors for ubiquitylation and downregulation. These data provide a framework for understanding how variants in MEGF8 cause the inherited multiple-congenital-disorder Carpenter’s syndrome (CRPTS) and sporadic cases of heterotaxy with severe congenital heart defects^10,18,24^.

## Results

### Structure of the MMM complex

Structural studies of membrane protein complexes containing large, multi-domain single-pass TM proteins like MEGF8 (**Figure S1A)** are challenging due to low expression yield, flexible linkages between domains, and compositional heterogeneity. We combined several technical strategies to generate an MMM sample amenable to structure determination by cryo-EM. First, we used a minimal MEGF8 construct (MEGF8^ΔR1R2^) that can ubiquitylate SMO and attenuate HH signaling^10,12^. MEGF8^ΔR1R2^ lacks a major portion of its ECD containing two pseudo repeats (R1 and R2), but retains an extracellular and juxtamembrane Stem domain required for its interaction with MOSMO^12^ (**Figures 1A and 1B**, and **Figure S1A**). To abolish auto-ubiquitylation and increase expression, we mutated a conserved linchpin residue in the MGRN1 RING domain (Arg317 in human or Arg318 in mouse). This residue is known to mediate key interactions with E2-Ub in other RING proteins^25–27^ and is essential for attenuating HH signaling in cells^23^, but is otherwise dispensable for the integrity of the RING fold (**Figures 1B and S1B**). Finally, to increase the size of the MMM complex, we developed a single-chain camelid antibody (nanobody NB270) that binds to both the MEGF8-MOSMO and MMM complexes with nanomolar affinity. The MMM components were co-expressed in human cells, solubilized in detergent micelles, and purified to homogeneity by tandem affinity purification followed by size-exclusion chromatography (SEC) (**Figures S1C and S1D**). The presence of MOSMO stabilized MEGF8^ΔR1R2^, consistent with a previous observation that the abundance of MEGF8 is reduced in *Mosmo^−/−^* cells^12^ (**Figure S1E)**. Expression of MGRN1 did not significantly impact the abundance of the MEGF8^ΔR1R2^-MOSMO complex (**Figure S1F**).

A monodisperse sample of the MMM-NB270 complex in glycol-diosgenin (GDN) detergent micelles (**Figure S1C)** was applied to cryo-EM grids and its structure determined to an average resolution of 3.03 Å (**Figure S2A, Table S1, Video S1,** and **Supplementary note 1**). A focused refinement of the extracellular and TM regions of MEGF8-MOSMO and NB270 improved the resolution of the MEGF8-MOSMO complex, yielding a 2.65 Å map (**Figure 1C** and **Table S1**). We did not observe clear cryo-EM density for the intracellular portion of MEGF8 (residues 2615-2778) nor for MGRN1, perhaps because MGRN1 was bound to MEGF8 via a flexible linkage (**Figure S2A**). We also determined the cryo-EM structure of the MMM complex bound to a different nanobody, which revealed a similar architecture but also lacked interpretable density for MGRN1 (**Figure S2B** and **Supplementary note 1**). AlphaFold (AF) modeling predicted a cytoplasmic solitary α-helix (SAH) in MEGF8 that seamlessly extends from the hydrophobic TM helix to the MGRN1-interacting ICD^28^ (**Figure 1B**). To reduce conformational heterogeneity and increase the resolution of the MEGF8 ICD and MGRN1, we replaced the SAH with a rigidified ER/K α-helix^29^ (MEGF8^ΔR1R2^-HelixMut, **Figures S1A and S1D**) and determined the cryo-EM structure of the helix-stabilized MMM complex to 3.29 Å overall resolution (**Figures 1D, 1E, and S2C, Video S1,** and **Supplementary note 1**). This structure revealed additional, intracellular density for the MEGF8 ICD and MGRN1. Even though the intracellular map is at a lower resolution compared to the MEGF8 TM and Stem domains, we were able to place and refine the MEGF8 ICD-MGRN1 complex using an AlphaFold (AF) model (**Figure 1E**).

MEGF8, MOSMO and MGRN1 bind in a 1:1:1 stoichiometry (**Figure 1E**). The extended TM-SAH helix of MEGF8 links MOSMO and MGRN1 through distinct interactions on opposite sides of the membrane. On the extracellular side, the β-sandwich Stem domain is perched like a cap atop the 4-TM helix bundle of MOSMO, making extensive contacts with its two extracellular segments (ECS1 and ECS2, **Figures 1B and 1E**) that build a compact 5 stranded β-sheet. The Stem domain has a short linker to the MEGF8 TM helix, which continues for ∼35 Å into the cytoplasm as a SAH (**Figure 1E**). MEGF8 ICD and MGRN1 form a unique complex where the polypeptide chains from both partners are intertwined to form a single globular structural unit. The MEGF8 ICD makes contacts with the MGRN1 RING domain and positions it beneath the inner leaflet of the plasma membrane (**Figure 1E** and **Video S1**).

### A membrane-embedded structural platform formed by the MEGF8-MOSMO interaction

Evolutionary sequence analysis placed MOSMO (previously called ATTHOG) within a sub-clade of tetraspans that includes the claudin-like proteins, best known as components of intercellular tight junctions^11^. Our structural data support this assignment: the four TM helices of MOSMO (α1 to α4) form a left-handed bundle crowned on the extracellular face by a β-sheet domain, a feature of the claudin fold^30,31^ (**Figures 2A, S3A, and S3B**). While the TM domains of MOSMO and claudins are similar, the extracellular antiparallel β-sheet domain of MOSMO, formed by β1-4 strands from ECS1 and β5 strand from ECS2, is divergent, likely because of its unique role in mediating an interaction with the MEGF8 Stem domain (**Figures 2A, S3A and S3B, Supplementary note 2**). A conserved disulfide bond between β1 and β2 in ECS1 of MOSMO and an extensive network of hydrogen bonds with water molecules and between side chains in ECS1 and ECS2 form a structured platform for interaction with MEGF8 (**Figures 2B and 2C, Video S1**).

**Figure 2.**
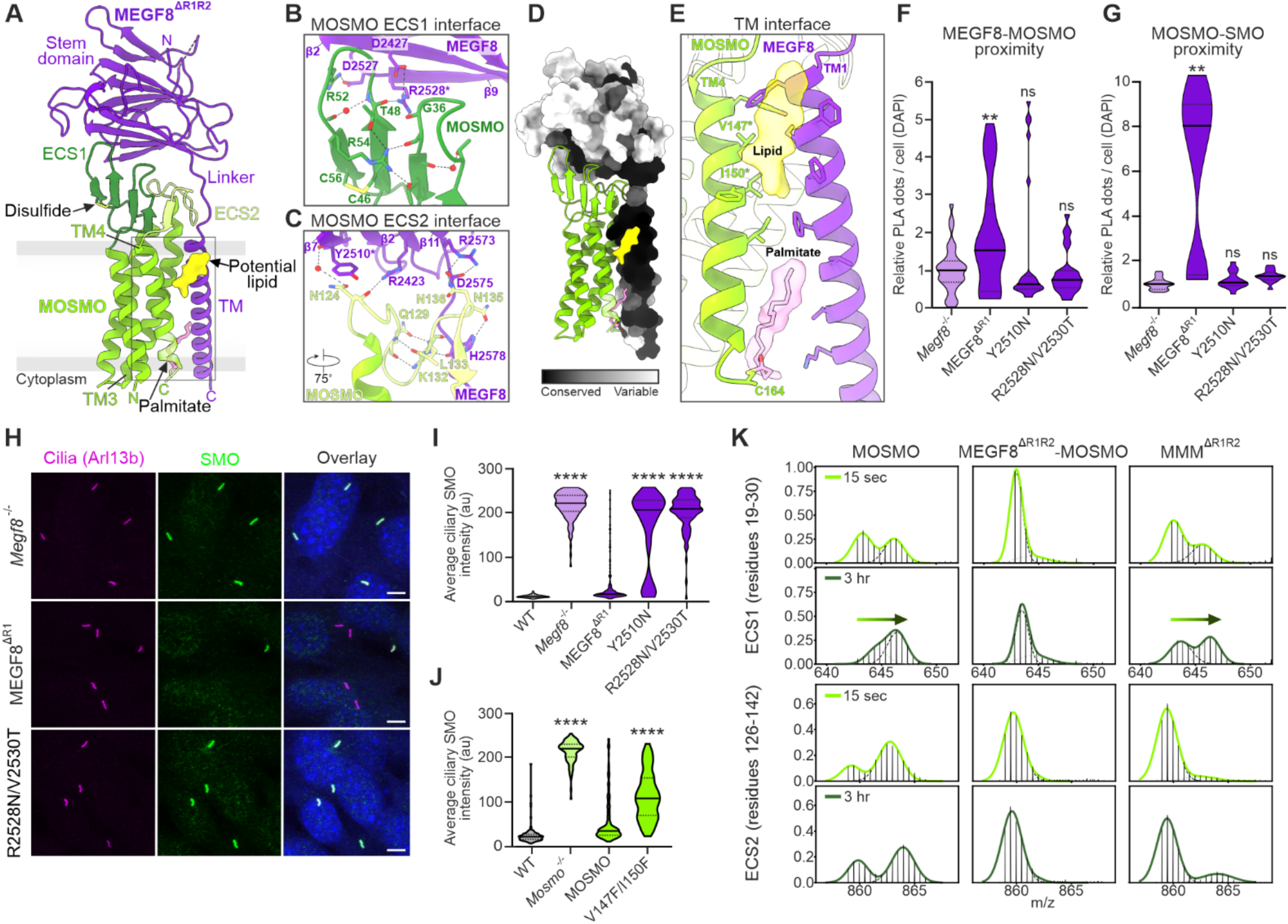
MOSMO forms extensive contacts with the MEGF8 Stem and TM domains. (**A**) Cartoon representations of the 2.65 Å MEGF8^ΔR1R2^-MOSMO complex structure showing the extracellular interface between the MEGF8 Stem domain and MOSMO ECS1/2, and the intramembrane interface between the single MEGF8 TM and MOSMO TM3 and TM4. Color coding is as in Figure 1. (**B** and **C**) Close-up views of the interfaces between MOSMO ECS1/2 and the MEGF8 Stem domain. Residues are depicted in stick representation with water as red spheres and hydrogen bonds depicted as dotted lines. (**D**) The solvent accessible surface representation of MEGF8^ΔR1R2^ with mapped sequence conservation reveals the highest conservation at the TM interface with MOSMO. (**E**) The cryo-EM maps for the palmitate covalently linked to MOSMO Cys164 in the inner leaflet (Figure S4, A to E) and potential sterol ligand in the outer leaflet. (**F** and **G**) Proximity between the indicated components measured by PLA (relative to *Megf8*^−/−^ cells) in *Megf8*^−/−^ NIH/3T3 cells stably expressing WT MEGF8ΔR1 or MEGF8 carrying mutations (Y2510N or R2528/V2530T) in the Stem domain. See Figure S5A for validation of PLA. (**H** to **J**) Representative images (H) and quantitation (I and J) of SMO abundance at primary cilia (marked by Arl13b) in *Megf8^−/−^* NIH/3T3 cells stably expressing MEGF8^ΔR1^ variants (H and I) or *Mosmo^−/−^* cells expressing MOSMO variants (J). Figure S5, B and C show protein abundances. (**K**) HDX-MS spectra and bimodal Gaussian fits for MOSMO peptides from ECS1 (top) and ECS2 (bottom) in various complexes at 15 s and 3 hours after D2O addition (3-and 15-min time points are shown in **Figures S7B and S7C**). Violin plots summarize *n*=15 images (each with 100 cells) (F and G) or *n*∼100 cilia (I and J), and show the median (solid line) and interquartile range (dashed lines). *p*-values (ns>0.05, **p<0.01, ****p < 0.0001) were determined using one-way ANOVA (each cell line compared to *Megf8^−/−^* cells) (F and G), or the Kruskal-Wallis (with Dunn) test (each mutant cell line compared to add back of the corresponding WT protein) (I and J). See also **Figures S3 to S7**.

MEGF8 and MOSMO form extensive, evolutionarily conserved interactions between their extracellular and TM domains (**Figure 2D**). The MEGF8 Stem domain buries a large surface area of 953 Å^2^ against the MOSMO ECD, with a shape complementarity score of 0.7, indicating a biologically relevant assembly^32–34^. The MEGF8-MOSMO extracellular interface is formed by 11 hydrogen bonds between the MEGF8 Stem domain, the MEGF8 TM linker, and the extracellular loops of MOSMO (**Figures 2B and 2C)**. The Stem domain adopts a β-sandwich fold with a core of nine β-strands arranged into two twisted sheets with complex strand connectivity (**Figure 2A, Figures S3C-S3F, Supplementary note 3**). One of these sheets is draped atop MOSMO and extends three arginine fingers (Arg2423 in β2, Arg2528 in β9, and Arg2573 in β11 strand) to make contacts with the MOSMO ECS1 and ECS2 (**Figures 2B and 2C**). Salt bridges between MEGF8 Asp2427 (β2) and Arg2528 (β9), as well as between Asp2575 and Arg2573 (β11), anchor the arginine fingers at the extracellular interface (**Figures 2B and 2C**, and **Video S1**). The MEGF8-Stem domain is connected to the single TM helix by a short linker of 7 predominantly polar residues, supported by hydrogen bonds between MEGF8-His2578 and the MOSMO-ECS2 Asn135 and Leu133 (**Figure 2C**). These interactions stabilize the ECS2 loop, which is typically unresolved in other claudin family proteins ^35,36^ (**Figures 2C and S3B, Supplementary note 3**).

Within the membrane, the interaction between the single MEGF8 TM helix and TM helices 3 and 4 from MOSMO includes two lipid moieties (**Figure 2E**). A palmitoyl chain covalently attached to the conserved Cys164 in the short cytoplasmic tail of MOSMO^11^, confirmed by mass spectrometry and atomistic molecular dynamics (MD) simulations (**Figures S4A-S4E, Table S2**), is sandwiched at this interface in the inner leaflet of the membrane. The palmitoyl group remained stably bound in simulations within a lipid bilayer, where it blocked bulk membrane lipids from accessing the pocket (**Figures S4D and S4E**). A density resembling a putative sterol is also found in the outer leaflet (**Figures S4F and S4G**). Although we modelled a GDN sterol moiety, the density does not unambiguously identify the lipid molecule (**Figures S4F and S4G, Video S1**).

Our structural data led us to test the functional relevance of the MEGF8-MOSMO interaction, which has remained enigmatic. While the birth defect phenotypes of *Megf8^−/−^* and *Mosmo^−/−^* mice are very similar, suggesting a shared function, MOSMO is not strictly required for MEGF8-MGRN1 to ubiquitylate SMO in a cell-based assay where the proteins are transiently over-expressed^10,12^. Structure-guided mutations at the MEGF8-MOSMO interface were introduced into MEGF8^ΔR1^, since the full-length MEGF8 cDNA is too large to clone and express in cells. Transgenes coding for variants were stably expressed in HH-responsive *Megf8^−/−^* NIH/3T3 cells. For low, near-endogenous expression, we inserted the transgenes into a single, defined genomic locus using the Flp-In recombinase system^37^. For higher-level expression, useful when the variants were less stable, we used lentiviral delivery^10^.

To assess interactions between MMM components and SMO, which cannot be detected by immunoprecipitation, we turned to proximity ligation assays (PLA) in fixed cells. PLA is a sensitive interaction assay that can detect proximity (<40 nm) between two proteins in their native cellular context without detergent solubilization or immunoprecipitation^38^. PLA detected proximity between SMO and each of the three MMM components in cells (**Figure S5A**). Introduction of ectopic N-glycosylation sites at the MEGF8 Stem interface with either ECS1 (MEGF8 R2528N/V2530T) or ECS2 (MEGF8 Y2510N) reduced MEGF8-MOSMO proximity, showing that these mutations destabilized their interaction and thus confirming the observed interface in our cryo-EM structures (**Figure 2F**). These interface mutations also reduced the PLA proximity signal between MOSMO and SMO (**Figure 2G**).

As an *in vivo* assay for MMM activity, we leveraged the observation that loss of any of the MMM components results in the massive accumulation of its direct substrate SMO in primary cilia in both cultured NIH/3T3 cells and mouse embryos, thereby enhancing sensitivity to HH ligands^10,12^ (**Figure 2H**). Ciliary SMO abundance, measured by quantitative immunofluorescence microscopy, can be used as a sensitive and graded measure of MMM activity in intact cells^10^. Ectopic N-linked glycosylation sites at the MEGF8-MOSMO interface abolished the ability of the MMM complex to attenuate SMO abundance at primary cilia (**Figures 2H, 2I, and S5B**). Mutations that introduced bulky side chains in MOSMO TM4 (V147F/I150F) at the interface with the putative sterol and MEGF8 (**Figure 2E**) reduced MMM activity in cells (**Figures 2J and S5C**). However, mutation of MOSMO Cys164 to Ser to prevent palmitoylation did not change MMM activity in our expression system (**Figures S5D and S5E**). We conclude that the MEGF8-MOSMO interaction is required for this E3 complex to both recognize and ubiquitylate its substrate receptor in cells.

### Local conformational dynamics of the MEGF8-MOSMO complex

The lower resolution of MGRN1 and MEGF8 ICD in our cryo-EM structure of the MMM complex led us to hypothesize that the MMM complex may exhibit conformational fluctuations that facilitate its function. To identify these, we carried out HDX-MS (**Table S3**). First, we compared HDX in MOSMO alone to its exchange profile in the presence of MEGF8. The vast majority of peptides exhibit enhanced protection (slower exchange) in the presence of MEGF8, consistent with stabilization of the extensive MEGF8-MOSMO binding interface observed in our cryo-EM structures (**Figures S6A and S6B**). The same regions of MOSMO also showed protection in the trimeric MMM complex across all time points (**Figures S6C and S6D)**.

Intriguingly, HDX-MS revealed that MGRN1 binding can modulate the dynamics at the MEGF8-MOSMO interface. In MOSMO alone, peptides show bimodal mass-isotope distributions in ECS1 and 2 (**Figures 2K and S7A-S7C**). Such bimodal behavior can arise when the rate of interconversion between exchange-competent and exchange-incompetent conformations is slower than the rate of deuterium labeling^39^. The addition of MEGF8 to MOSMO suppresses MOSMO dynamics, such that ECS1 and ECS2 remain fully protected from exchange. Addition of MGRN1 to MEGF8-MOSMO restores the presence of the second, exchange-competent state, but slows the conversion between these populations (**Figures 2K and S7A-S7C**). These data suggest that MGRN1, which makes no direct contacts with MOSMO in our structure, is allosterically coupled to the extracellular MEGF8-MOSMO interface. The enhanced conformational flexibility of ECS1 and 2 in the MMM complex may reflect orientational heterogeneity of the MEGF8 SAH when bound to MGRN1 (**Figures 1C and 1E**).

### A hand-in-glove interaction between the MEGF8 ICD and MGRN1

Sequence analysis and inspection of our structure shows that MGRN1 is composed of four segments: an N-terminal pre-engager domain, an engager domain (also called DAR2, **Figures S3H and S3I**, **Supplementary note 4**), a RING domain (responsible for E2-Ub recruitment and activation), and an intrinsically disordered region spanning the C-terminal 200 amino acid residues (**Figure 3A**)^2,11,40^. RING-family ligases often have separate substrate or scaffold recognition domains that are flexibly linked to the catalytic RING domain^41,42^. Unexpectedly, our cryo-EM structure showed that MGRN1 binds the MEGF8 ICD through a multi-partite interaction, using the pre-engager, engager and RING domains to incorporate MEGF8 into the E3 machinery (**Figures 3A and 3B, Video S1**). Deletion of the pre-engager, engager or RING domains compromised both the interaction between MEGF8 and MGRN1 in a co-immunoprecipitation assay (**Figure S8A**) and the ability of the MMM complex to attenuate ciliary SMO abundance in cells (**Figure S8B**). The disordered segment (not visible in the structure) is dispensable for the MEGF8-MGRN1 interaction but is required for SMO regulation (**Figures S8A and S8B)**. To cleanly assay the function of MGRN1 variants without interference from co-expressed wild-type proteins, we stably expressed them in *Mgrn1^−/−^;Rnf157^−/−^* double null cells lacking both MGRN1 and RNF157, a closely related and partially redundant E3 ligase^10,11^.

**Figure 3.**
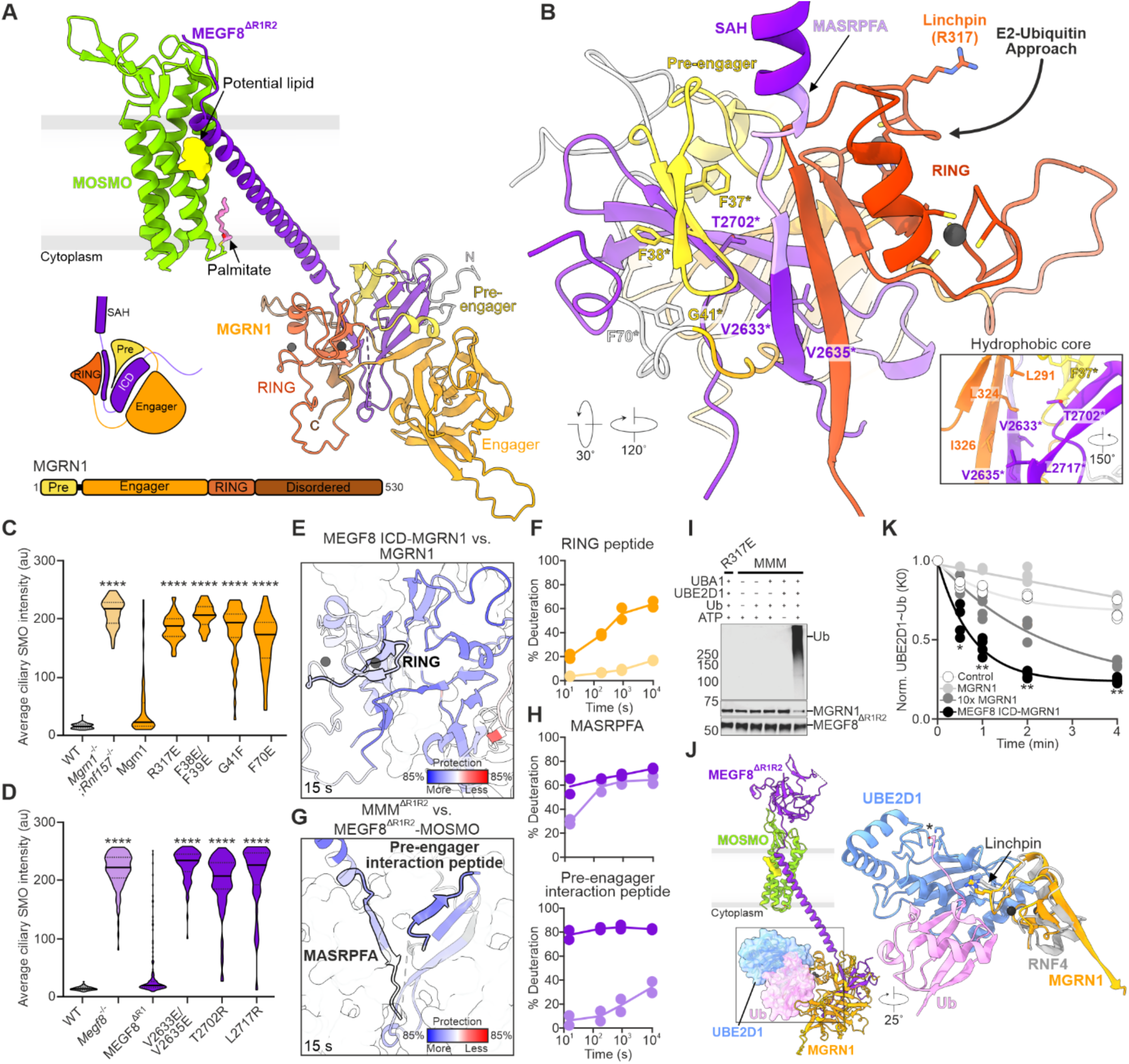
A multi-partite interaction between MEGF8 and MGRN1. (**A**) Cartoon representation of the MMM complex, showing how the MGRN1 pre-engager (Pre), engager, and RING domains surround the MEGF8 ICD (schematized in the inset). Zn^2+^ ions bound in the RING domain are shown as gray spheres. (**B**) View of the MEGF8 ICD-MGRN1 interface, with an inset showing a close-up of the hydrophobic core formed by side chain interactions of both proteins (in stick representation). The critical MASRPFA peptide (light purple segment in MEGF8) forms a flexible linker that allows the MEGF8 ICD-MGRN1 structural unit to swivel around the MEGF8 SAH (Figure S10). The MGRN1 R317 linchpin residue points away from the complex and marks the site of E2-Ub engagement. Residues targeted for structure-guided mutagenesis and functional analysis are highlighted with asterisks (C and D). (**C** and **D**) SMO abundance in primary cilia of *Mgrn1^−/−^* ; *Rnf157^−/−^* NIH/3T3 cells expressing MGRN1 variants (C) or *Megf8^−/−^* cells expressing MEGF8^ΔR1^ variants (D). **Figures S8E and S8F** show protein abundances. (**E** and **F**) Cartoon representation of MGRN1 colored by the percent difference in deuterium exchange in the MEGF8 ICD-MGRN1 complex relative to MGRN1 alone (E). Example of an uptake plot showing percent deuteration over time for a peptide in the RING domain (F, residues 283-296, MGRN1 in dark orange, MEGF8 ICD-MGRN1 complex in light orange). Figure S9D shows uptake plots of peptides in the MGRN1 pre-engager and engager domains. (**G** and **H**) Cartoon representation of the MEGF8 intracellular region colored by the percent difference in deuterium exchange at 15 s in the MMM complex compared to the MEGF8-MOSMO complex (G). Examples of uptake plots (H) for the MASRPFA peptide (a.a. 2625-2643) (upper graph) and a peptide that binds to the engager domain (lower graph). MEGF8-MOSMO: dark purple, MEGF8-MOSMO-MGRN1: light purple. (**I**) Auto-ubiquitylation assay to assess catalytic activity of the MMM complex used for cryo-EM (with or without the MGRN1 R317E linchpin mutation). (**J**) Superposition of the RNF4 (E3)-UBE2D1 (E2)-Ub complex (PDB ID: 4AP4^26^) onto the MGRN1 RING domain in our MMM complex structure, with UBE2D1 (blue) and Ub (pink) shown as solvent-accessible surfaces (left). A close-up of the structural alignment in cartoon representation (right) shows the relative position of the linchpin Arg resides in RNF4 and MGRN1 in stick representation. An amide bond (asterisk) between UBE2D1-K85 and Ub-G76 was used as a stable mimic of the labile thioester linkage. RMSD: 1.162 Å for 33 equivalent Cα positions across the RNF4 and MGRN1 RING domains. (**K**) Decay of the UBE2D1-Ub(K0) conjugate as a function of time after the addition of 40 mM lysine alone (control) or lysine and the indicated E3 enzymes (*n*=4 per time point, lines are best-fit single exponential curves). MGRN1 alone was used at the same concentration or 10-fold (10X) higher concentration compared to the MEGF8 ICD-MGRN1 complex. Figure S11E shows a representative gel. Violin plots summarize *n*∼100 cilia and show the median (solid line) and interquartile range (dashed lines) (C and D). *p*-values (*p < 0.05, **p < 0.01, ****p < 0.0001) were determined using the Kruskal-Wallis (with Dunn) test (each mutant cell line compared to the corresponding WT add back in C and D, and the E3 complex compared to MGRN1 alone in K). F and H show mean (+/−SD, *n*=2) values. See also **Figure S8 to S11**.

One side of the MEGF8-MGRN1 interface is formed by a MEGF8 β-strand that immediately follows the SAH and is inserted into the center of the MEGF8 ICD-MGRN1 structure (**Figure 3B**). This MEGF8 β-strand (residues 2630-2636) extends a β-sheet through interaction with an edge β-strand (residues 322-331) of the RING domain. Consequently, the RING domain is rigidly attached to the MEGF8 ICD, orienting it such that the MGRN1 linchpin residue is pointing away from MEGF8 and hence free to engage the E2-Ub conjugate and catalyze Ub transfer to substrate receptors (**Figure 3B**). This structural role of the RING is notable because in most E3s, including other membrane-embedded E3s, the RING domain is flexibly tethered to the other parts of the structure and is dedicated to Ub transfer^42–44^.

Despite this atypical structural role, the MGRN1 RING domain is a typical RING domain containing a cross-braced structure of two zinc fingers (**Figure 3B**). Cys and His residues coordinate the two zinc ions and intervening loops contain the residues, such as the Arg317 linchpin, critical for E2-Ub engagement. A charge-reversal mutation in the linchpin residue (R317E, used in our cryo-EM sample) abolished ligase activity (as measured by ciliary SMO abundance) without disturbing the association of MEGF8 and MGRN1 (**Figure S8A and S8B**). Other residues critical for MGRN1 RING domain catalytic function also abrogated MMM activity in cells without reducing association with MEGF8^10^ (**Figure S1B)**. In the PLA assay, SMO-MGRN1 proximity required integrity of the RING domain, suggesting a role in substrate recognition (**Figures S8C and S8D**).

On the side opposite the RING, a three-stranded β-sheet from the MEGF8 ICD is sandwiched by a β-strand from the MGRN1 pre-engager domain on one side and an edge β-strand from the engager domain on the other, resulting in an extended anti-parallel β-sheet (**Figure 3A**). The pre-engager domain inserts an antiparallel β-hairpin like a wedge into the space between the central β-strand and 3-stranded β-sheet of the MEGF8 ICD (**Figures 3A and 3B**). This intertwined mode of recognition is quite distinct from other interactions that recruit soluble cytoplasmic proteins to the membrane, suggesting that the structural code of MGRN1-class E3 ligases (which also include RNF157 and CGRRF1) for recognition of a TM receptor like MEGF8 is likely to be complicated and only apparent after accommodation to the MGRN1 scaffold. In contrast, SH2, SH3 or PDZ domain-mediated interactions that recruit proteins to cytoplasmic tails of receptor tyrosine kinase or membrane-localized scaffolds involve the recognition of disordered peptide motifs by a globular domain^45^.

Structure-guided mutagenesis and functional testing in *Mgrn1^−/−^;Rnf157^−/−^* NIH/3T3 cells^10^ using ciliary SMO abundance assays validated the importance of several contacts between MGRN1 and the MEGF8 ICD. Mutations in the pre-engager (F38E/F39E, G41F) and the engager (F71E) domains of MGRN1 impaired activity to the same extent as the R318E linchpin mutation (**Figures 3C and S8E**). In the MEGF8 ICD, mutations at the interfaces with the RING (V2633E/V2635E) or engager (T2702R, L2717R) also abolished activity (**Figures 3D and S8F**).

### Stability of the MEGF8 ICD-MGRN1 interaction

To complement our cryo-EM structural analysis, we carried out HDX-MS of MGRN1 alone and compared it to HDX-MS of a complex between MGRN1 bound to the MEGF8 ICD alone or the complete MEGF8-MOSMO complex (**Figures 3E, 3F, and S9A-S9D**). MGRN1 peptides from the RING, pre-engager, and engager domains showed marked protection from hydrogen-deuterium exchange when present in a complex with MEGF8, even at time points as long as 3 hours, supporting the formation of a stable complex (**Figures 3E, 3F, and S9A-S9D**). To study conformational dynamics of the MEGF8 ICD, we compared the binary MEGF8-MOSMO complex to the ternary MMM complex, since MEGF8 could not be purified in isolation (**Figures 3G, 3H**, **and S9E-S9G**). The regions of the MEGF8 ICD that make contacts with MGRN1 in our MMM complex cryo-EM structure show extensive protection from deuterium exchange, while in the absence of MGRN1 these same regions show near maximal (80-90%) exchange even at short time points (**Figures 3G, 3H**, **and S9E-S9G**), suggesting that MGRN1 binding imposes structural order on the MEGF8 ICD. Enhanced protection of the cytoplasmic SAH from deuterium exchange suggests that its helical structure is stabilized in the ternary complex containing MGRN1 (**Figure S9G**).

A notable feature of the MEGF8 ICD-MGRN1 interaction is the short loop that connects the MEGF8 SAH to the first β-strand of the ICD (**Figure 3B**). This loop is formed by a peptide sequence (MASRPFA) that has been long noted to be highly conserved in MEGF8, ATRN, ATRNL1 and related proteins across evolution^10,22,46^. Since deletion of the MASRPFA motif abrogates MMM activity, it was proposed to be the critical sequence motif recognized by MGRN1^10^. Surprisingly, our cryo-EM structure shows that the MASRPFA motif does not play a major role in recognition but rather properly links the MEGF8 SAH to the globular ICD-MGRN1 unit (**Figure 3B**). In atomistic simulations, this motif exhibited substantial conformational flexibility (**Figures S10A and S10B**, with the MEGF8 ICD-MGRN1 unit able to swivel freely around this point. Consistent with these MD simulations, the MASRPFA peptide was more dynamic than other peptides at the MEGF8 ICD-MGRN1 interface in HDX experiments, showing increased protection from deuterium exchange in the presence of MGRN1 at 15 s but not at later time points (**Figure 3H**). Since the MGRN1 RING is held rigidly, we speculate that the MASRPFA provides the mobility required for the multi-ubiquitylation of receptor substrates^10^.

### RING activation by the MEGF8 ICD-MGRN1 interaction

To assess whether the MEGF8 ICD-MGRN1 complex seen in our structure was competent for Ub transfer, we developed an *in vitro* system with purified proteins to measure ubiquitylation **(Figure 3I)**. Using the MEGF8^ΔR1R2^-MOSMO-MGRN1 complex used for cryo-EM (but without the inactivating R317E linchpin mutation), we screened 34 E2s and identified UBE2D family E2s as the most effective at supporting auto-ubiquitylation (**Figure S11A**). Auto-ubiquitylation was specific because it depended on E1, E2 and ATP and was prevented by the R317E linchpin mutation (**Figure 3I**), which also abrogates regulation of SMO and HH signaling in cells^23^ (**Figure S8B)**. Modeling the interaction between the MGRN1 RING and an E2-Ub conjugate, using a crystal structure of a primed RNF4 RING-E2-Ub complex^26^, revealed that an E2-Ub can be accommodated in the MMM complex without steric clashes with the membrane or any other domains (**Figure 3J** and **Video S1**). Importantly, the linchpin residues of MGRN1 and RNF4 occupy similar positions relative to the labile E2-Ub thioester bond, suggesting a shared mechanism for E2-Ub activation^25–27^.

To directly test if MGRN1 or the MEGF8 ICD-MGRN1 complex can activate its cognate E2, we measured the rate of Ub discharge from a pre-formed conjugate formed between UBE2D1 and lysine-less Ub (UBE2D1-Ub(K0)) to an excess of the small nucleophile lysine in solution^47,48^. This assay allows the isolated assessment of the ability of an E3 to activate an E2 in the absence of substrates. MGRN1 alone and the MEGF8 ICD-MGRN1 complex (**Figure S11B**, both expressed and purified from bacteria) were capable of auto-ubiquitylation in the presence of UBA1 (E1), UBE2D1 (E2), Ub and ATP. However, the MEGF8 ICD-MGRN1 complex was significantly more active: 10-fold higher concentrations of MGRN1 were required to transfer the same amount of Ub into poly-Ub chains (as seen by free Ub depletion) (**Figures S11C and S11D**). Consistent with this observation, the rate of Ub(K0) discharge from a UBE2D1-Ub(K0) conjugate was much faster in the presence of the MEGF8 ICD-MGRN1 complex compared to MGRN1 alone (**Figures 3K and S11E**). These results suggest that the interaction between the MEGF8-ICD and MGRN1 stabilizes the active conformation of the MGRN1 RING domain (in addition to optimally orienting it for Ub transfer to receptor substrates, **Figure 3B**).

### Molecular basis for disease-causing mutations in MEGF8

Disruption of genes encoding any of the three MMM components causes mid-gestational lethality in mouse embryos with defects affecting the heart, skeleton, limb and brain. Most notably, embryos lacking MMM components suffer from heterotaxy, a defect in left-right patterning of the body axis that leads to severe congenital heart defects incompatible with life^10,12^. A very similar spectrum of birth defects is seen in a subset of human patients with CRPTS, an autosomal-recessive multiple-congenital-anomaly syndrome, that carry mutations in the *MEGF8* gene^18,24^. These patients suffer from left-right patterning defects, complex congenital heart defects, polysyndactyly and craniosynostosis (**Figure 4A**). Pathogenic variants in MMM genes have also been discovered in human babies suffering from sporadic heterotaxy and congenital heart defects^10^. While some of the birth defects, such as polydactyly, are almost certainly caused by elevated HH signaling^12^, it is likely that other signaling pathways are also affected in these patients^49^.

**Figure 4.**
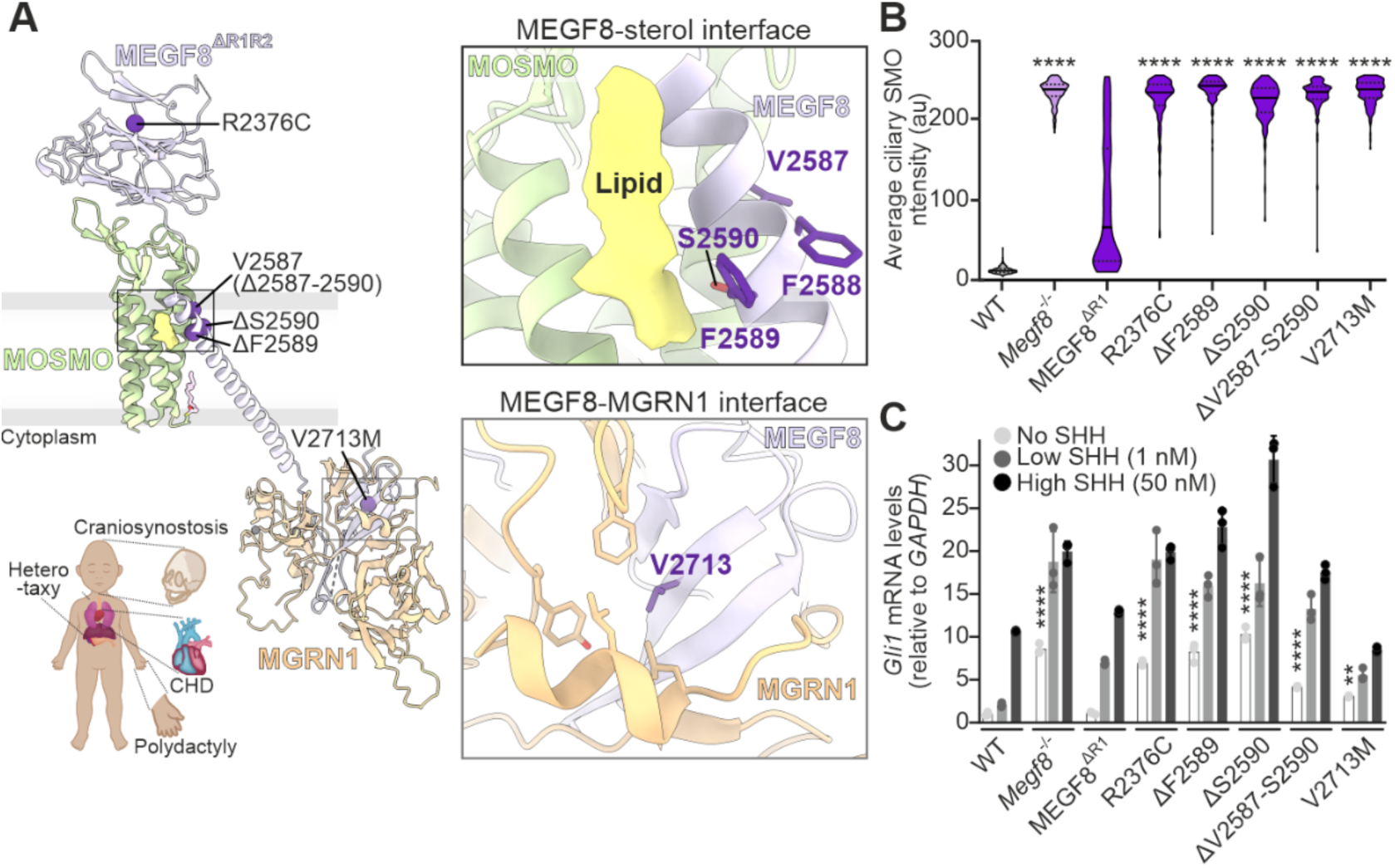
MEGF8 mutations found in Carpenter’s Syndrome patients enhance sensitivity to HH ligands. (**A**) Cartoon representation (left) of the MMM complex with mapped residues altered in Carpenter’s syndrome patients (depicted as Cα positions in spheres). Close-up views (right) of selected residues in stick representation at the MEGF8-MOSMO (top) and MEGF8-MGRN1 (bottom) interfaces. The inset highlights key phenotypic features of the syndrome. (**B** and **C**) SMO abundance (B) and SHH ligand sensitivity (C) in *Megf8^−/−^* NIH/3T3 cells expressing the MEGF8^ΔR1^ variants found in Carpenter’s syndrome depicted in A. Figure S12 shows protein abundances of the variants and an independent experiment using a different expression system that yields higher overall protein abundances. The abundance of *Gli1* mRNA, a metric of signaling strength encoded by a direct HH target gene, was measured by qRT-PCR after exposure to the indicated SHH concentrations (C). Violin plots summarize *n*∼100 cilia and show the median (solid line) and interquartile range (dashed lines) (B). *p*-values (****p < 0.0001) were determined using the Kruskal-Wallis (with Dunn) test (each mutant cell line compared to the corresponding WT add back) (B). Bars show the mean (+/− SD, n=3) and *p*-values (**p < 0.01, ****p < 0.0001) determined using one-way ANOVA (with Dunnett) test (no SHH condition in each mutant cell line compared to no SHH condition in WT add back) (C). **See also Figure S12.**

Our structure of the MMM complex allowed us to investigate the biochemical consequences of missense mutations identified by human genetic analysis of CRPTS patients and families^24^. We focused on mutations in residues that were resolved in our structure, including one (R2376C) in the MEGF8 Stem, one in the MEGF8 ICD (V2713M) predicted to disrupt hydrophobic interactions between MEGF8 and MGRN1, and two mutations at the interface where the MEGF8 TM helix contacts MOSMO (**Figure 4A**). All mutants could be stably expressed in *Megf8^−/−^* NIH/3T3 cells, though some (e.g. V2713M) were present at lower abundances (**Figure S12A**). These MEGF8 mutants were also expressed using lentiviral transduction to ensure that observed phenotypes were not caused only by lower protein abundances (**Figures S12B and S12C**). MMM complexes containing the MEGF8 variants mapped in CRPTS patients were compromised in their ability to clear SMO from primary cilia (**Figure 4B)** and to suppress HH target gene transcription (**Figure 4C**). Cells carrying these human *MEGF8* disease variants showed elevated basal HH signaling and were hyper-sensitive to the ligand Sonic Hedgehog (SHH), achieving similar levels of target gene transcription at 50-fold lower SHH concentrations (**Figure 4C**). This markedly higher sensitivity is likely to disrupt the development of tissues (like the limb) that depend on exposure of cells to HH ligand concentrations that are precisely calibrated in space and time.

### An extended α-helix as a central architectural element

A striking feature of the MMM complex is the central TM-SAH helical element of MEGF8 that links MGRN1 in the cytoplasm to extracellular and TM domains of MOSMO (**Figure 1E**). While the structures and relative positions of MOSMO and the MEGF8 Stem domain are similar in the wild-type and the ER/K helix-stabilized MMM complex, the orientation of the TM-SAH is different. The ER/K stabilized helix pivots inward (towards MOSMO) by 22° around the fulcrum formed by its point of contact with MOSMO (**Figure 5A and Video S1**). Consequently, MGRN1, which is suspended at the end of this helical lever arm in the cytoplasm, is in a different position relative to the membrane-embedded TM-Stem platform. Interestingly, MEGF8^ΔR1R2^ could clear SMO from cilia when expressed in *Megf8^−/−^* cells, whereas the ER/K stabilized helix construct was inactive (**Figures 5A and 5B**). Notably, both MEGF8 variants are expressed and associate normally with MGRN1 and MOSMO, as shown by SEC (**Figures S1C and S1D**).

**Figure 5.**
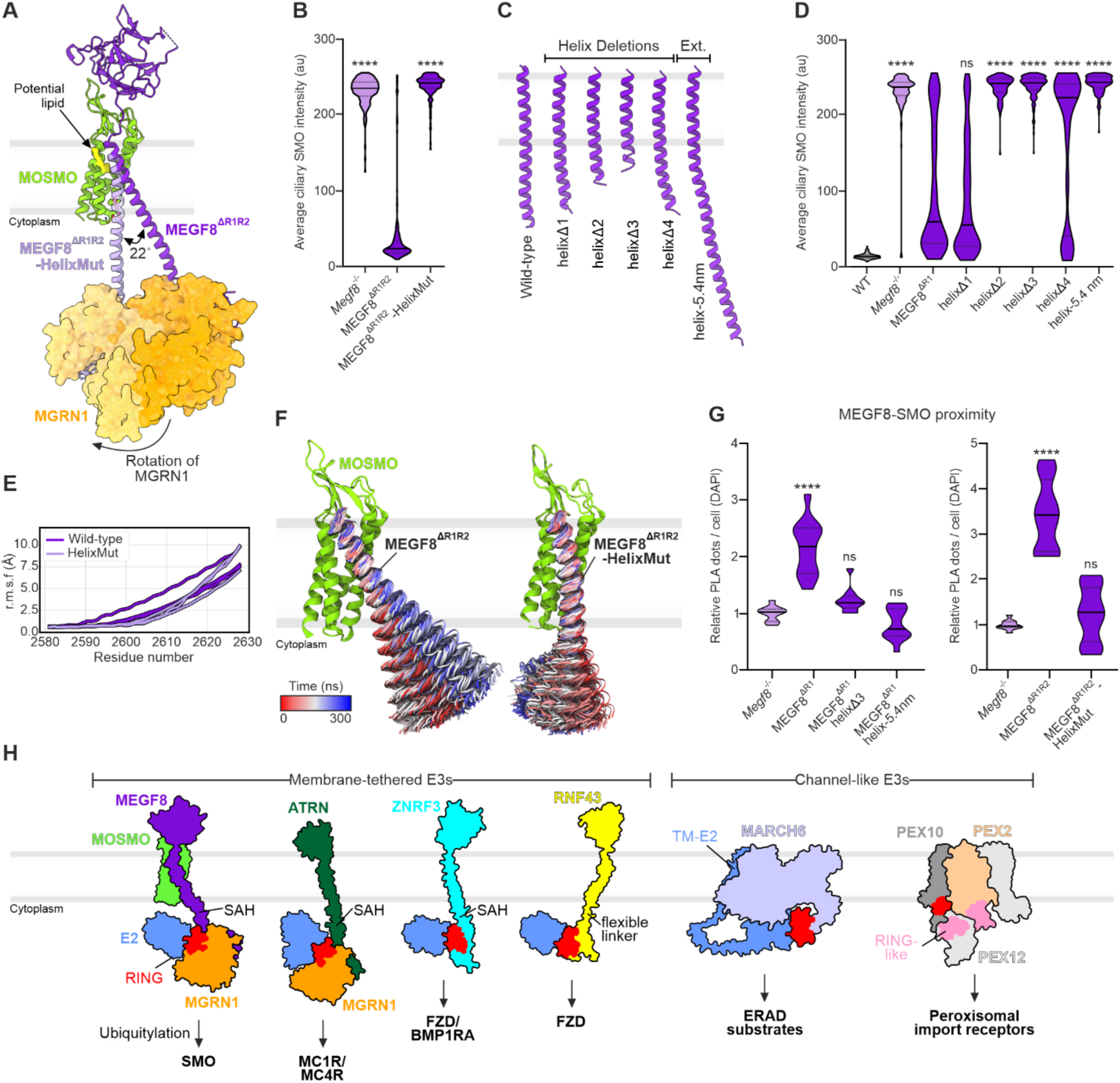
The length and flexibility of a central extended alpha helix is a key design element for MMM function. (**A**) Superimposition of MMM and helix-stabilized MMM complexes based on MOSMO (RMSD: 0.386 Å for 152 equivalent Cα positions). MGRN1 is shown as a solvent accessible surface. The stabilizing ER/K helix in the MEGF8^ΔR1R2^-HelixMut (see Figure S1A for sequences) occludes the palmitate binding site. (**B**) Ciliary SMO abundances in *Megf8*^−/−^ NIH/3T3 cells expressing either MEGF8^ΔR1R2^ or MEGF8^ΔR1R2^-HelixMut. (**C** and **D**) Impact of various mutations that change the length of the MEGF8 SAH on the ability of MEGF8^ΔR1^ to reduce the abundance of SMO at primary cilia (D). See Figure S13A-S13E for protein abundances of variants, data from independently generated cell lines and other controls. (**E)** Root mean square fluctuation (r.m.s.f.) of the MEGF8 TM-SAH region after fitting to MOSMO. Calculations were performed across 6 independent 300 ns simulations of the WT (dark purple) and HelixMut (light purple) constructs. (**F**) The MEGF8 SAH shows substantial conformational flexibility in MD simulations. 300 evenly distributed structures of the native and ER/K stabilized MEGF8 TM-SAH are colored according to simulation time. (**G**) Proximity between endogenous SMO and the indicated MEGF8 variants in NIH/3T3 cells was measured by PLA as described in Figure 2F. (**H**) Surface presentations for various membrane-spanning E3s highlight the position of a canonical RING domain (shown in red in all structures) relative to the inner leaflet of the membrane. Proteins are drawn to scale by tracing around the solvent-accessible surface of structures or structural models. The presence of a SAH spacer and ubiquitylation substrates are noted below. AF3 models were used for Attractin (ATRN)-MGRN1, ZNRF3, RNF43 and MARCH6; cryo-EM structures for the MMM (this study) and PEX (PDB ID: 7T92^50^) complexes. Violin plots summarize *n*∼100 cilia (B and D) and *n*=10 images (each with 100 cells) (G), and show the median (solid line) and interquartile range (dashed lines). *p*-values (ns>0.05, ****p < 0.0001) were determined using the Kruskal-Wallis (with Dunn) test (each mutant cell line compared to the corresponding WT add back) (B and D) and one-way ANOVA (each cell line compared to *Megf8^−/−^* cells) (G). See also **Figures S13 and S14**.

Together with our structural analysis, these data suggest a model where the helix functions as a tether to anchor MGRN1 at an optimal distance below the inner leaflet to recognize and ubiquitylate the cytoplasmic surface of SMO. To test this idea, we shortened the cytoplasmic SAH segment, carefully maintaining the helical register and avoiding removal of the key MASRPFA residues (**Figure 5C)**. While smaller deletions of ∼1 turn of the helix (helixΔ1 and Δ4) were able to partially clear SMO from cilia, longer deletions resulted in complete loss of function (**Figures 5D and S13A-S13E**). An insertion predicted to extend the helix by ∼5.4 nm (“helix-5.4nm”) also abrogated activity (**Figures 5C and 5D**). The importance of the helix is also supported by the prior observation that a catalytically active MEGF8 ICD-MGRN1 complex simply dangling from the inner leaflet of the plasma membrane failed to ubiquitylate SMO ^10^.

To investigate the conformational landscape of the TM-SAH, we carried out six independent 300 ns atomistic simulations of the MMM complex with either the wild-type helix or the stabilized ER/K variant embedded in a 1-palmitoyl-2-oleoyl-sn-glycero-3-phosphocholine (POPC) bilayer. These simulations revealed that the SAH has substantial conformational flexibility. The TM-SAH segment behaves more like a flexible tether than a fixed ruler, occupying a conformational space that resembles a cone with its apex formed by the contact point between the helix and MOSMO (**Figures 5E, 5F**, **and S13F-S13H**). The conformational fluctuation is smallest where the MEGF8 TM contacts MOSMO and progressively increases through the membrane and into the cytoplasmic SAH (**Figure 5E**). The wild-type SAH, but not the ER/K variant, is capable of bending away from the MOSMO platform and interacting with the lipid bilayer, potentially allowing it to reach a receptor embedded in the membrane **(Figure S13H and Video S2**). Consistent with this model, PLA assays demonstrated that both changes in helix length or orientation impaired the ability of the MMM complex to recognize SMO in cells (**Figure 5G**).

These data highlight the importance of helix orientation in MMM function, which is a consequence of the unusual angle at which the helix traverses the membrane distal to its contact point with MOSMO (**Figure 5A**). This interface is a site of three pathogenic CRPTS mutations (**Figure 4A**). In summary, our data support the model that the continuous MEGF8 TM-SAH, which makes no direct contacts with either MOSMO or MGRN1, organizes the overall geometry of the complex to enable ubiquitylation of receptor substrates embedded in the plasma membrane.

## Discussion

Amongst the ∼600 RING-type E3 ligases, ∼50 have been classified as TM ligases by the criteria that they contain predicted TM helices in their polypeptide chains^2^. While many remain unstudied, TM E3s have been implicated in protein quality control and protein transport at organelles. Available structures of TM E3s include two involved in ER-associated protein degradation (HRD1 and DOA10/MARCH 6) and the PEX E3 complex involved in protein transport across the peroxisomal membrane^43,44,50,51^. These assemblies are characterized by the formation of a channel in the membrane for the recognition of unfolded polypeptide substrates. The RING domain is flexibly attached to the rest of the ligase and positioned just below the membrane channel to enable ubiquitylation concomitant with polypeptide transport (**Figure 5H**).

Compared to these channel-like E3s, the MMM complex presents a fundamentally different architecture adapted to its task of recognizing and ubiquitylating folded signaling receptors on their cytoplasmic surfaces (**Figure 5H**). Key features include (a) the use of a flexible SAH domain to suspend the catalytic RING at a specific depth and orientation below the inner leaflet of the plasma membrane, (b) use of MOSMO to provide a table-like β-sheet for MEGF8 Stem recognition and a 4 TM helical anchor in the membrane, (c) stabilization and orientation of a canonical RING domain through an intricate structural interaction. These principles imply that simple proximity of the MGRN1 RING to the membrane is insufficient for receptor ubiquitylation; rather it must be presented in a specific position and orientation relative to the receptor substrate. These MMM features are likely to be shared with other membrane-spanning E3 ligases that target cell surface receptors (**Figures 5H and S14A**). AF models for the ATRN-MGRN1 complex and ZNRF3, a single TM E3 ligase that targets Frizzled receptors for WNT ligands^5^, predict the presence of an SAH that functions as a flexible spacer between the inner leaflet of the membrane and the RING domain (**Figure 5H**). Strikingly, in each of these cases, the models predict that the RING domain is held via a rigid linkage using a β-sheet extension motif (**Figure S14A**), as we observe for MGRN1 in the MMM complex structure (**Figure 3B**). The dramatic protection of RING peptides from deuterium exchange when MGRN1 is bound to MEGF8 (**Figures 3E and 3F**) supports the notion of RING stabilization within the complex.

Modeling the MMM complex with SMO and an E2-Ub conjugate shows that MGRN1 is positioned by these interactions in a pose that would allow it to engage E2-Ub and activate it for transfer to a lysine-rich patch on the cytoplasmic surface of SMO or other receptor substrates **(Figures S14B-S14E)**. Overall, the MMM complex resembles a fishing rig designed to suspend bait at a specific distance below the water’s surface: MOSMO serves as a float in the membrane that uses the central MEGF8 helix as a flexible tether to suspend MGRN1 at a distance and orientation optimal to capture and ubiquitylate SMO.

### Limitations of this study

There are several outstanding questions for future research, including how SMO is recognized selectively, the identity of other receptor substrates (suggested by the widespread birth defect phenotypes), and the role of the massive pseudo-repeated MEGF8 ECD (modeled by AF as two tandem globular domains emanating from the Stem)^28^ **(Figure S14B)**. While this ECD is not required for SMO recognition and ubiquitylation^10^, it may allow MMM regulation by luminal or extracellular ligands, akin to how the TM E3 ligases for WNT receptors (ZNRF3 and RNF43) are regulated by R-Spondin ligands^3^ **(Figure 5H)**. Finally, the number of membrane-spanning ubiquitin ligases like the MMM complex remains uncertain since the non-covalent association of an E3 with a membrane protein scaffold is difficult to predict by sequence alone. However, these architecturally unique assemblies are likely optimized for recognition of receptor substrates, making them useful partners for therapeutic protein targeting approaches against mutant or disease-driving receptors^52^.

## Acknowledgements

We thank Thomas Walter and Karl Harlos of The Division of Structural Biology for technical support, Helen Duyvesteyn of Oxford Particle Imaging Center for assistance with electron microscopes, Sophie Shoemaker and Shawn Costello for sharing their expertise in HDX MS analysis and technical support during MS experiments, and Eva Beke and Irene Tiku for technical support during Nanobody discovery. Work was supported by Cancer Research UK (C20724/A26752 and DRCRPG-May23/100002 to C.S.) and the National Institutes of Health (GM118082 and HL157103 to R.R). C.W. was supported by a Wellcome Trust DPhil studentship (218482/Z/19/Z). The Wellcome Centre for Human Genetics, Oxford, is funded by Wellcome Trust Core Award (090532/Z/09/Z). Oxford Particle Imaging Center was funded by a Wellcome Trust JIF award (060208/Z/00/Z) and is supported by equipment grants from Wellcome Trust (093305/Z/10/Z). Research in S.L.R. group is supported by UKRI Future Leaders Fellowship (MR/Y01975X/1) and Wellcome Trust Discovery Award (301619/Z/23/Z). Research in the laboratory of J.H.K. was supported by NIH/NIGMS (GM132518), NIH/NCI (CA015704), and Dick and Anne Schneider. L.M.N. was supported by the Walter V. And Idun Berry Postdoctoral Fellowship. N.L. and S.M were supported by NIH funding (K99GM148823 and R35GM149319, respectively). S.M. is a Chan Zuckerberg Biohub Investigator. This work benefited from access to the Nanobodies4Instruct Centre (VID40422 PID23766), we acknowledge the support and use of resources of Instruct-ERIC, part of the European Strategy Forum on Research Infrastructures (ESFRI) and the Research Foundation - Flanders (FWO) for their support to Nanobody discovery.

## Author contributions

Conception: C.W., L.M.N., J.H.K., C.S. and R.R. Design of study: C.W., L.M.N., G.H., P.P., E.P., N.L., G.V.P., L.M., J.S., S.M., J.F.B., S.L.R., L.C., J.H.K., C.S. and R.R. Data collection: C.W., L.M.N., G.H., P.P., E.P., D.L., N.L., L.G.., R.C. and J.H.K. Data analysis: C.W., L.M.N., G.H., P.P., E.P., D.L., N.L., G.V.P., L.G., L.M., S.M., S.L.R., L.C., J.H.K., C.S. and R.R. Supervision: C.S. and R.R. Acquisition of funding: S.M., S.L.R., J.H.K., C.S. and R.R. Writing and editing: C.W., L.M.N.,G.H., C.S. and R.R., with input from all authors.

## Supplemental information

**Document S1.** Figures S1-S14, Tables S1-S3, STAR methods, supplementary notes and supplementary references.

**Video S1.** Structural overview of the MMM complex, related to Figures 1-4.

**Video S2.** Molecular dynamics simulations of the MEGF8 SAH, related to Figure 5.

**Video S3.** Molecular dynamics simulations of the MEGF8-SMO transmembrane interface, related to Figure S14.

## Supplementary figures

**Figure S1:**
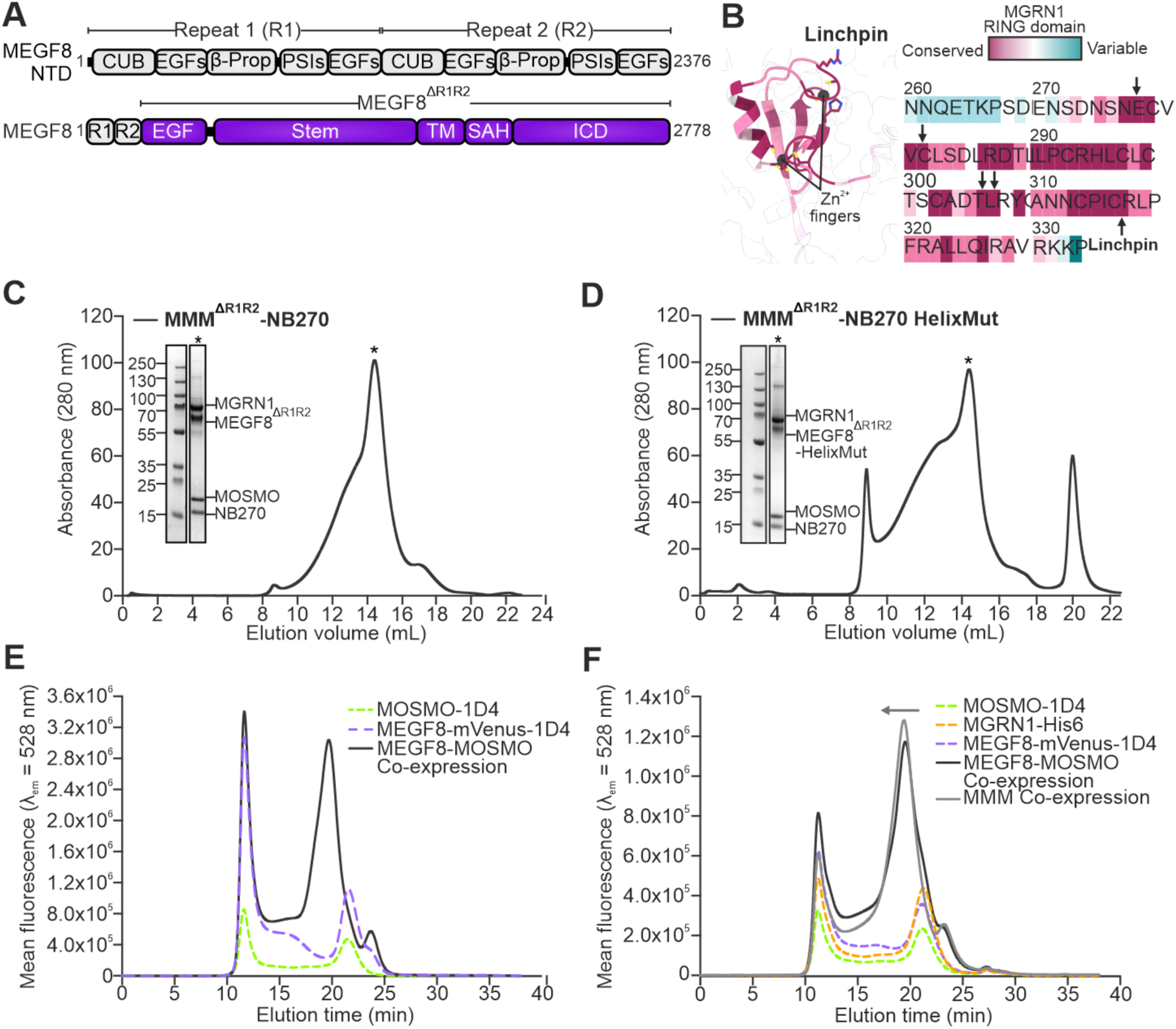
Expression and purification of MMM complexes. (**A**) Domain organization of full-length MEGF8, including the two large multi-domain pseudo repeats (R1 and R2) that were removed from our expression construct (MEGF8^ΔR1R2^). EGF, epidermal growth factor-like domain; TM, transmembrane helix; SAH, solitary α-helix; ICD, intracellular domain; CUB, <u>C</u>omplement protein C1r/C1s-<u>u</u>rEGF-<u>B</u>MP1 domain; β-prop, 6-bladed β-propeller domain; PSI, <u>P</u>lexin-<u>S</u>emaphorin-Integrin domain. (**B**) Conservation of the MGRN1 RING domain (based on 1500 sequences between 35-95% sequence similarity compared to human MGRN1) mapped onto an AlphaFold (AF) model of MGRN1. Arrows show the linchpin residues (R317), mutated to glutamate (R317E) in our expression construct, and other conserved RING domain residues that have been shown to be critical for regulation of SMO and HH signaling^10^. (**C** and **D)** Size-exclusion chromatography (SEC) profiles for wild-type and helix-stabilized MMM-NB270 complexes used for cryo-EM. Peak fractions (asterisk) were analyzed by reducing SDS-PAGE followed by Coomassie staining. (**E** and **F**) SEC analysis of MEGF8^ΔR1R2ICD^ fused to a fluorescent protein (mVenus) and co-expressed with either MOSMO alone or MOSMO and MGRN1. Mean fluorescence determined from two technical repeats. Arrow highlights a peak shift consistent with association of MGRN1 with the MEGF8^ΔR1R2ICD^-MOSMO complex (F).

**Figure S2:**
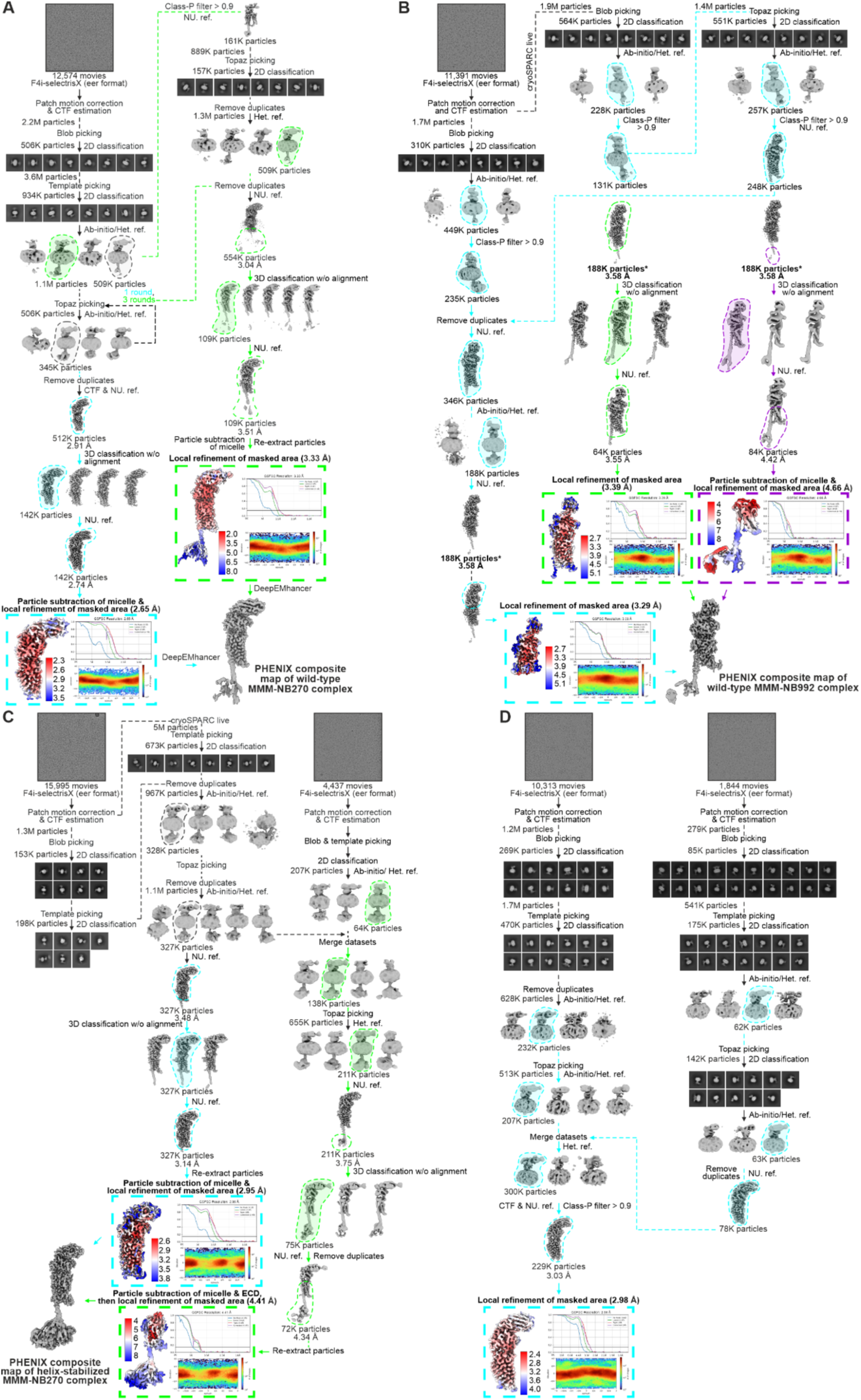
Cryo-EM data collection, processing and analysis scheme of our MMM and MEGF8-MOSMO complexes. Workflow for the image processing of the wild-type MEGF8^ΔR1R2^-MOSMO-MGRN1 complex with NB270 (A) and NB992 (B), the helix-stabilized MEGF8^ΔR1R2^-MOSMO-MGRN1-NB270 complex (C), and MEGF8^ΔR1R2ICD^-MOSMO-NB270 complex (D, without MGRN1 bound) in cryoSPARC v.4-2.1-4.6.2^53^. Representative micrographs for the MMM complexes embedded in vitreous ice are shown, followed by example 2D classes and the processing pipeline. Particles from two datasets collected on the same grid were merged for the final map (B and C). The Fourier shell correlation (FSC) curves indicating overall map resolution, local resolution maps and angular distributions of each map are shown in the final rectangle in each branch of the workflow. Local resolutions were computed using the cryoSPARC local resolution estimation tool. Composite maps were generated using locally refined maps in PHENIX. For the wild-type MMM-NB270 complex (A), maps were sharpened with DeepEMhancer^54^ using the highRes model, before generating a composite map in PHENIX^55^. See also **Supplementary note 1.**

**Figure S3:**
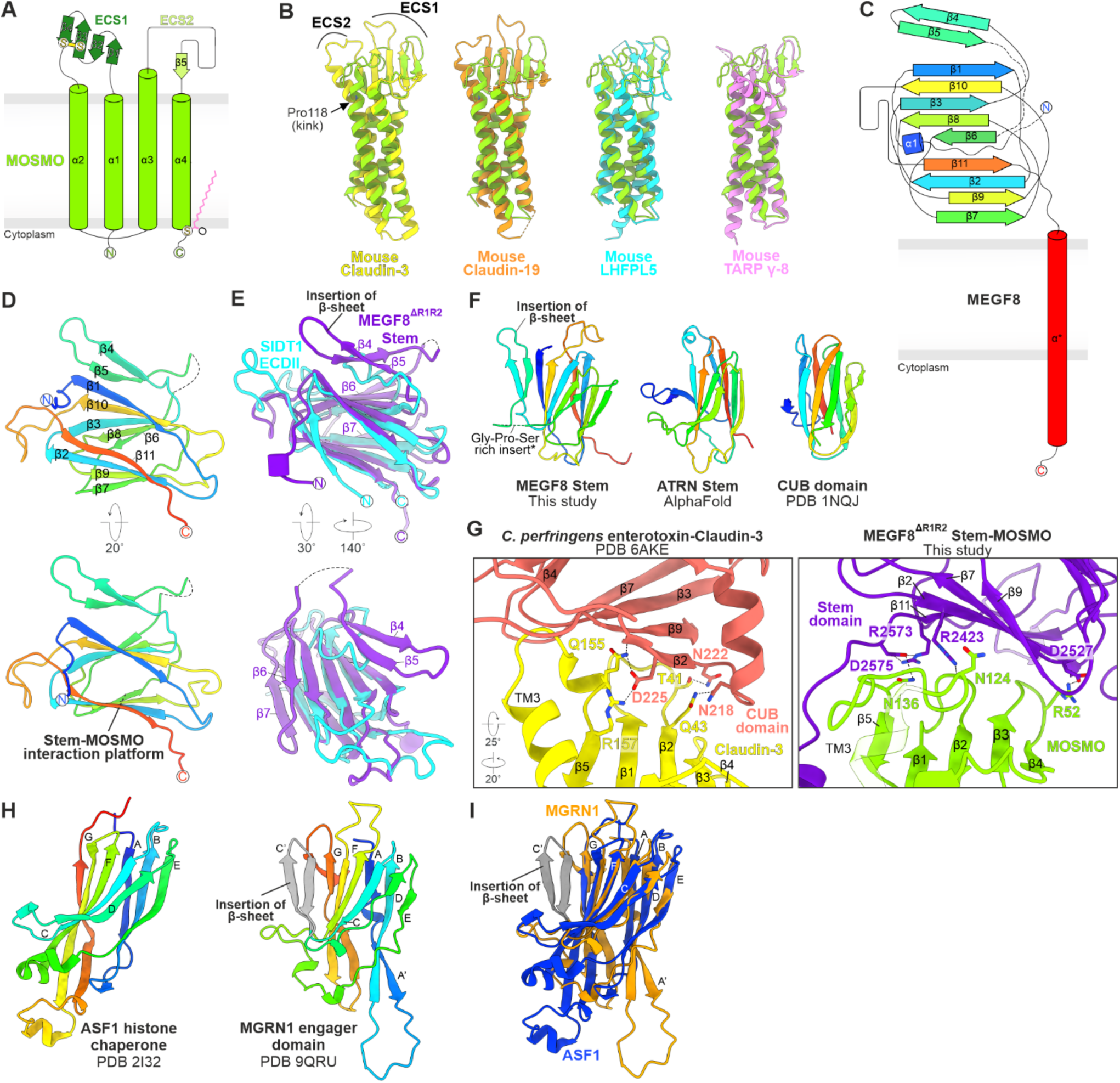
Analysis and comparison of the MOSMO, MEGF8 and MGRN1 structures. (**A**) Topology diagram of MOSMO, highlighting the five strands from two extracellular segments (ECS) that make up a β-sheet. A palmitate is covalently linked to C164 at the C-terminus. (**B**) Superposition of MOSMO (green) with crystal structures containing a claudin fold: mouse claudin-3 (yellow, PDB ID: 6AKF^56^, RMSD: 0.949 Å for 66 equivalent Cα positions), mouse claudin-19 (orange, PDB ID: 3X29^57^, RMSD: 1.139 Å for 76 equivalent Cα positions), mouse LHFPL5 (cyan, PDB ID: 6C14^35^, RMSD: 1.241 Å for 61 equivalent Cα positions), mouse TARP γ8 (pink, PDB ID: 7LEP^36^, RMSD: 0.875 Å for 42 equivalent Cα positions). A kink formed by Pro118 in the longer TM3 of MOSMO is highlighted by an arrow. (**C** and **D**) Organization of the β-sheets within the MEGF8 Stem domain, in rainbow coloring (from the N-terminus: blue to C-terminus: red). (**E**) Structural comparison of the MEGF8 Stem domain and human SIDT1 ECDII (PDB ID: 8JUL^58^). (**F**) MEGF8 and Attractin (ATRN) Stem domains share a CUB-like fold^59^. Structures are colored from N-terminus in blue to C-terminus in red. (**G**) The mouse Claudin-3 binding interface (yellow) with a C-terminal fragment of *Clostridium perfringens* enterotoxin (red) (PDB ID: 6AKE^56^, left panel) is similar to the analogous interface between MOSMO and the MEGF8 Stem domain (this study, right panel). (**H**) Ig-like topology of human histone chaperone ASF1 (PDB ID: 2I32^60^) and human MGRN1 engager domain (this study) colored with N-terminus in blue to C-terminus in red. MGRN1 contains an additional β-hairpin (C’, gray) inserted between the C and D β-sheets in the core fold. (**I**) Superimposition of ASF1 (PDB ID: 2I32^60^, blue) onto the MGRN1 engager domain (orange).

**Figure S4:**
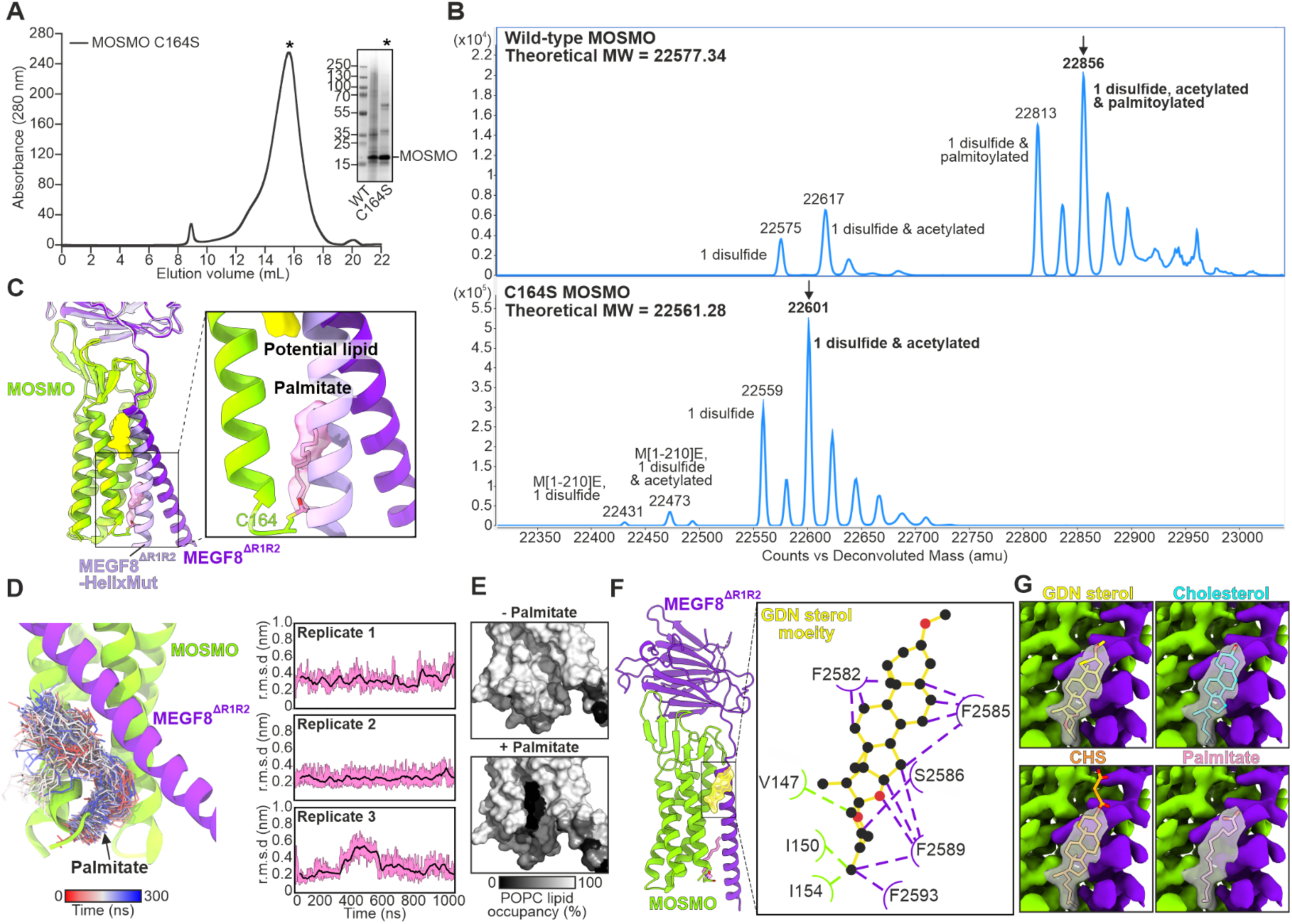
Analysis of putative lipid ligands at the intramembrane interface between MOSMO and MEGF8. (**A**) Representative SEC purification of a MOSMO mutant in which the Cys residue (C164) predicted to be palmitoylated ^11^ has been changed to a serine (C164S mutant). Peak fractions of wild-type and MOSMO-C164S were pooled and analyzed by reducing SDS-PAGE followed by Coomassie staining. (**B**) Denaturing intact mass spectrometry analysis of wild-type (top) and C164S MOSMO (bottom). For wild-type MOSMO, observed masses correspond to the theoretical mass of MOSMO with one disulfide bond (−2 Da, 100% occupancy), acetylation (+42 Da, 64 % occupancy) and palmitoylation (+238 Da, 75% occupancy). The peak representing palmitoylated MOSMO is indicated by an arrow in the top panel. For mutant MOSMO, observed masses correspond to the theoretical mass of the full-length MOSMO C164S mutant with one disulfide bond (−2 Da, 100 % occupancy), acetylation (+42 Da, 57% occupancy) and loss of the C-terminal lysine (−128 Da, 1% occupancy). (**C**) Superposition of the native MEGF8^ΔR1R2^-MOSMO (dark purple) and helix-stabilized MEGF8^ΔR1R2^-HelixMut-MOSMO (light purple) complexes based on MOSMO (RMSD: 0.386 Å for 152 equivalent Cα positions). Cryo-EM maps of the potential lipid and palmitoyl sites are shown in yellow and pink, respectively. The pocket containing the MOSMO C164-palmitoyl in the structure of MEGF8^ΔR1R2^-MOSMO complex is occluded in the MEGF8^ΔR1R2^-HelixMut-MOSMO complex in which the SAH is replaced by a rigidified ER/K helix. (**D**) The palmitoyl group remains stably bound over 3 × 1000 ns of atomistic MD simulation in a lipid membrane. 300 evenly distributed palmitoyl structures are colored according to simulation time. Plots (right) show the r.m.s.d. of the palmitoyl group after fitting to MOSMO. Raw data is shown in light pink, and the sliding average in black. (**E**) The palmitoyl group blocks membrane lipids from accessing the pocket in simulations. Surface representation of the protein is colored according to the per residue occupancy of 1-palmitoyl-2-oleoyl-sn-glycero-3-phosphocholine (POPC). (**F**) Putative lipid density (yellow) in the outer leaflet of the membrane from the cryo-EM map of wild-type MMM complex. Inset (middle) shows the residues from MOSMO (green) and MEGF8 (purple) that interact with a modeled glyco-diosgenin (GDN) detergent headgroup (in stick representation) as measured by LigPlot+^61^. (**G**) Comparison of different molecules placed into the cryo-EM map for the putative lipid shown in (F). CHS: cholesteryl hemisuccinate.

**Figure S5:**
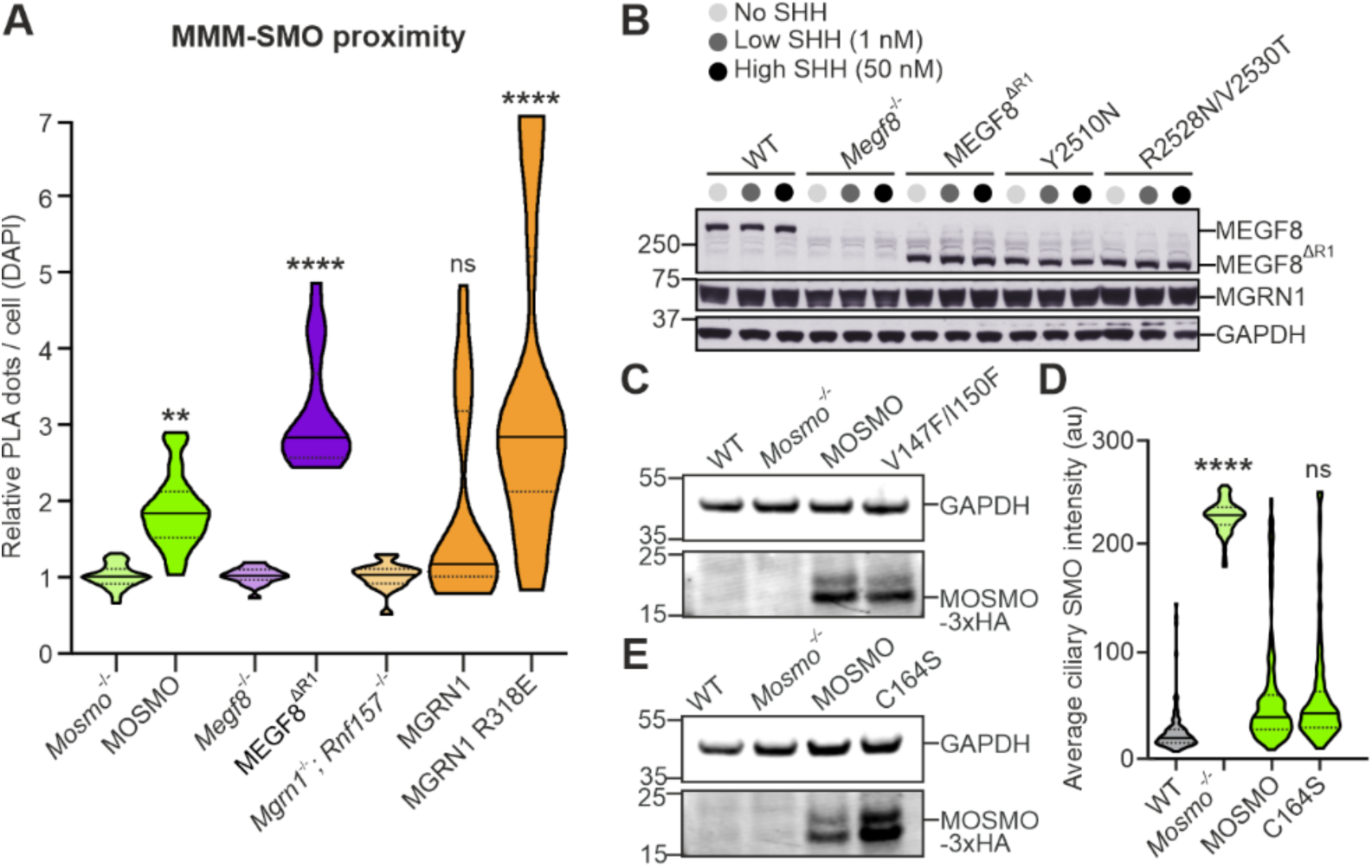
Structure-guided functional interrogation of the MEGF8-MOSMO interface. (**A**) Validation of the proximity ligation assay (PLA) used to assess proximity between endogenous SMO and each component of the MMM complex (related to Figures 2F, 2G, 5G, S8C, and S8D). MOSMO-3xHA, MEGF8^ΔR1^-1D4, and MGRN1-3xFLAG were stably expressed in NIH/3T3 cell lines (*Mosmo^−/−^*, *Megf8^−/−^*, *Mgrn1^−/−^;Rnf157^−/−^* respectively) in which the corresponding endogenous gene was disrupted. Data are plotted relative to the PLA signal measured in the corresponding knockout line to correct for non-specific signals. The higher signal seen with the inactive linchpin mutant of mouse MGRN1 (R318E, same position as R317E in the human protein) likely reflects a substrate-trapping effect. (**B** and **C**) Immunoblotting was used to measure the abundances of stably expressed variants of MEGF8^ΔR1^ (B) and MOSMO (C), corresponding to functional studies shown in Figure 2, I and J. (**D** and **E**) Ciliary SMO abundances in *Mosmo*^−/−^ cells stably expressing WT MOSMO or a mutant (C164S) that cannot be palmitoylated. Total protein abundances were assessed by immunoblotting (E). Violin plots summarize *n*=15 images (each with 100 cells) (A) and n∼100 cilia (D), and show the median (solid line) and interquartile range (dashed lines). *p*-values (ns>0.05, **p<0.01, ****p < 0.0001) were determined using one-way ANOVA (each cell line compared to its corresponding null control line) (A) or the Kruskal-Wallis (with Dunn) test (each mutant cell line compared to the MOSMO WT add back) (D).

**Figure S6:**
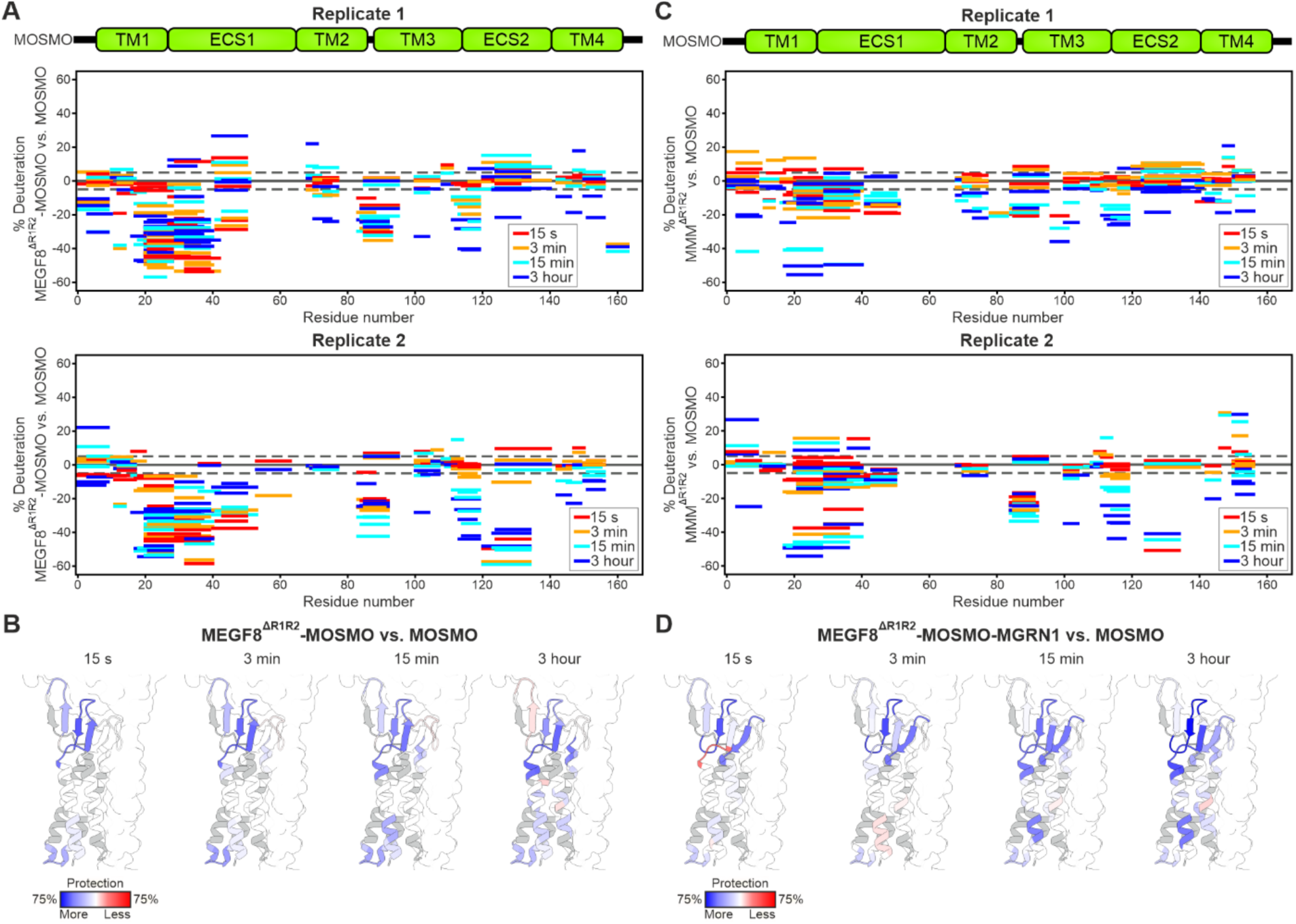
Analysis of hydrogen-deuterium exchange (HDX) rates of MOSMO peptides using mass spectrometry (MS). (**A**) Woods plots (2 replicates) show the difference in deuterium uptake in peptides from MOSMO when it is present in a complex with MEGF8^ΔR1R2^ compared to when present alone. Each bar is a peptide, with the bar length representing the peptide length, its position in the protein sequence indicated by the *x*-axis, and its color denoting the time point after deuterium addition at which the peptide was detected by MS. The position of the bar on the *y*-axis denotes the difference in percent deuteration of this peptide in the MEGF8^ΔR1R2^-MOSMO complex compared to MOSMO alone. Thus, negative *y*-values mean that a peptide was more protected from deuterium exchange in the complex. (**B**) Woods plot data from (A) was mapped onto a structure of the MOSMO in the MMM complex. Color scale shows the per residue difference in deuteration in the MEGF8^ΔR1R2^-MOSMO complex relative to MOSMO alone. (**C** and **D**) Woods plots and corresponding structural maps for MOSMO peptides when present in the trimeric MMM complex (with both MEGF8^ΔR1R2^ and MGRN1) compared to MOSMO alone. Data are represented as in (A and B).

**Figure S7:**
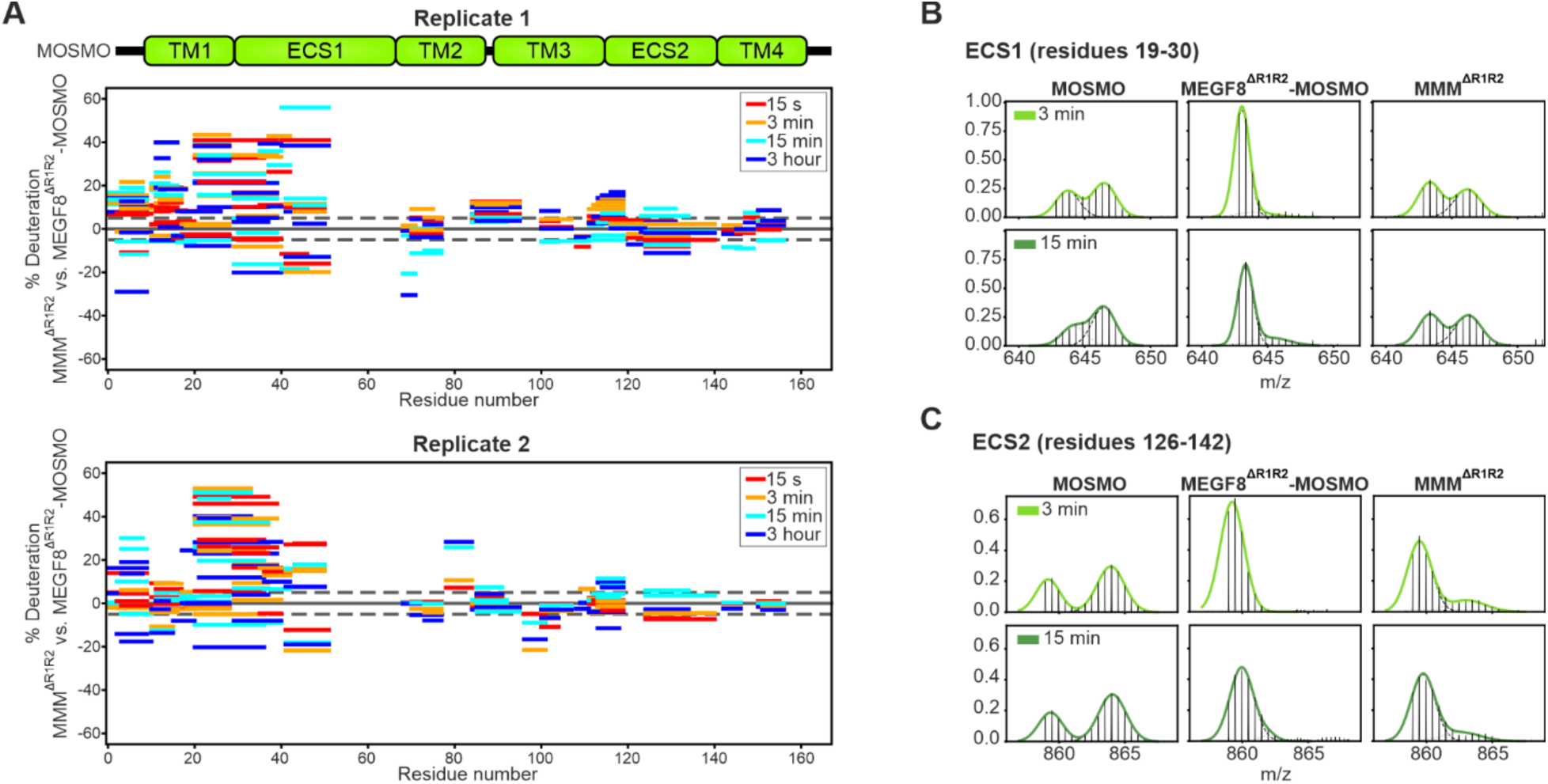
HDX-MS analysis of MOSMO peptides in the MMM complex. (**A**) Woods plots for MOSMO peptides from the MEGF8^ΔR1R2^-MOSMO-MGRN1 complex compared to the MEGF8^ΔR1R2^-MOSMO complex. Data are represented as described in Figure S6A. (**B** and **C**) HDX-MS spectra and bimodal Gaussian fits for MOSMO peptides from ECS1 (top) and ECS2 (bottom) in various complexes at 3 min and 15 min after D2O addition (15 s and 3-hour time points are shown in Figure 2K).

**Figure S8:**
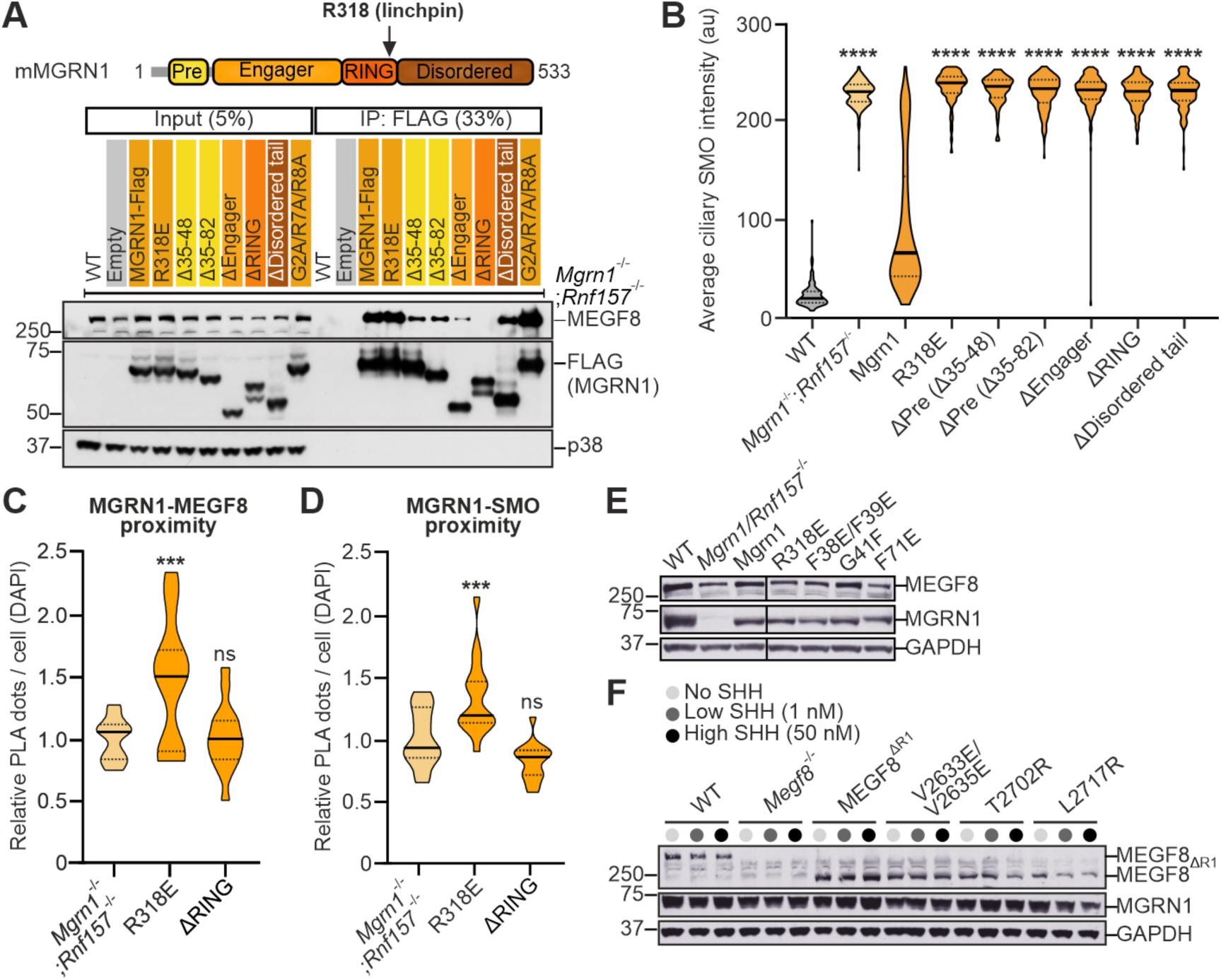
Structure-guided functional interrogation of the MEGF8-MGRN1 interface. (**A**) Immunoprecipitation (IP) followed by immunoblotting was used to measure the interaction between endogenous MEGF8 and various mutants of MGRN1, all appended with a C-terminal FLAG tag and stably expressed in *Mgrn1*^−/−^/Rnf157^−/−^ NIH/3T3 cells. 5% of the total input and 33% of the total IP were loaded on the gel. Domain composition of MGRN1 with amino acid boundaries is shown above to identify the position of the various point mutations and truncations. R318E is the equivalent RING linchpin mutation used for our cryo-EM construct in mouse MGRN1 (Figure 1); Δ35-48 and Δ35-82 are truncations within the pre-engager domain (Pre); ΔRING, ΔEngager and ΔDisordered are complete internal deletions of each domain; G2A/R7A/R8A is a triple point mutation that is predicted to abolish N-myristoylation of MGRN1. (**B**) Abundance of SMO at primary cilia in *Mgrn1*^−/−^/Rnf157^−/−^ NIH/3T3 cells stably expressing the MGRN1 variants shown in (**A**). (**C** and **D**) PLA was used to measure proximity between endogenous MEGF8 (C) or endogenous SMO (D) and the indicated variants of MGRN1-FLAG stably expressed in *Mgrn1*^−/−^/Rnf157^−/−^ NIH/3T3 cells. Data are represented relative to *Mgrn1*^−/−^/Rnf157^−/−^ cells. See Figure S5A for validation of PLA. (**E** and **F**) Immunoblotting was used to measure the abundances of MGRN1 variants (E) or MEGF8 variants (F) stably expressed in *Mgrn1*^−/−^/Rnf157^−/−^ or *Megf8*^−/−^ NIH/3T3 cells, respectively. These abundance controls correspond to the functional studies shown in Figures 3C and 3D. Violin plots summarize n∼100 cilia (B) or *n*=10 images (each with 100 cells) (C and D), and show the median (solid line) and interquartile range (dashed lines). *p*-values (ns>0.05, ***p<0.001, ****p < 0.0001) were determined using the Kruskal-Wallis (with Dunn) test (each mutant cell line compared to the MGRN1 WT add back) (B), and one-way ANOVA (each cell line compared to *Mgrn1*^−/−^/Rnf157^−/−^ cells) (C and D).

**Figure S9:**
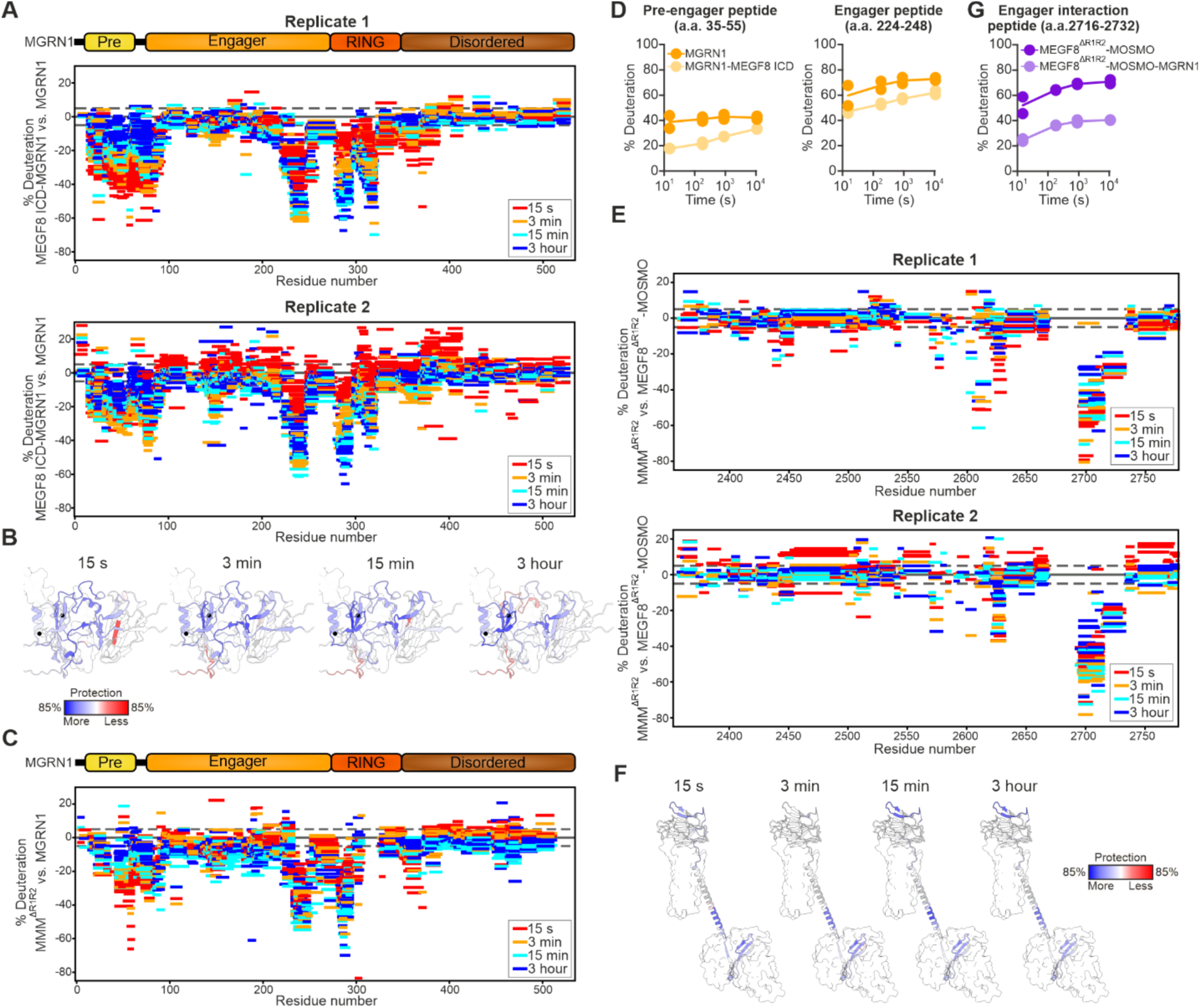
HDX-MS analysis of peptides involved in the MEGF8-MGRN1 interaction. (**A** to **C**) Woods plot for MGRN1 peptides in a MEGF8 ICD-MGRN1 complex (A, sample expressed in and purified from *E. coli*) or a MEGF8^ΔR1R2^-MOSMO-MGRN1 complex (C, our cryo-EM sample from Figure 1) compared to MGRN1 alone. Data are represented as described in Figure S6A. Woods plot data (**A**) was mapped onto the MGRN1 structure in cartoon representation with a color scale that shows the per residue difference in deuteration in the MEGF8 ICD-MGRN1 complex compared to MGRN1 alone (B). (**D**) Uptake plots show percent deuteration over time of peptides in the MGRN1 pre-engager (a.a. 35-55) and engager (a.a. 224-248) domains in MGRN1 or the MEGF8 ICD-MGRN1 complex. (**E** and **F**) Woods plots (E) and corresponding structural overlay (F) showing the difference in deuteration of MEGF8 peptides in the MEGF8^ΔR1R2^-MOSMO-MGRN1 complex compared to the MEGF8^ΔR1R2^-MOSMO complex. (**G**) Uptake plots of a MEGF8 ICD peptide (a.a. 2716-2732) that interacts with the engager domain of MGRN1.

**Figure S10:**
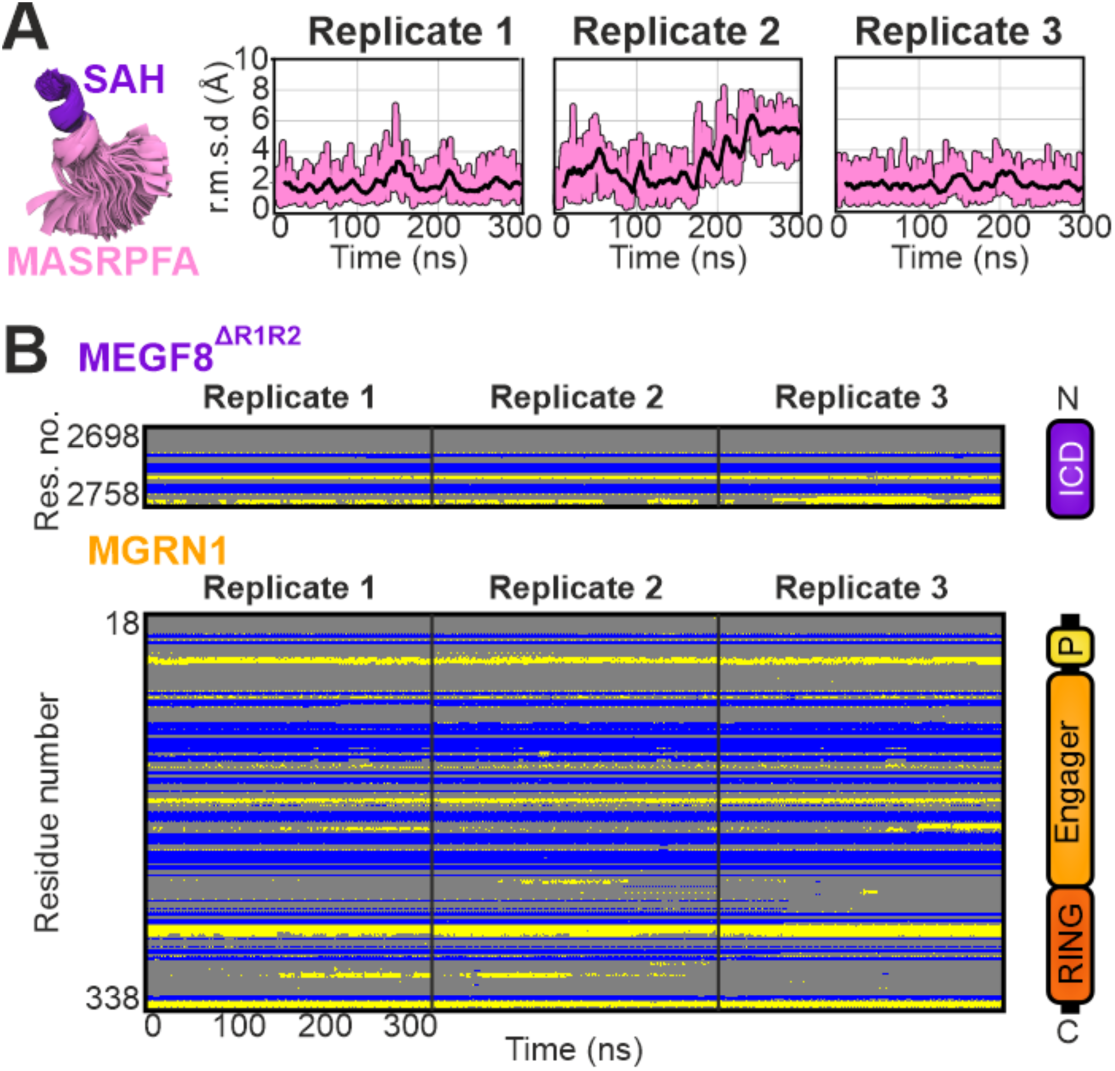
Molecular dynamics of the MEGF8 ICD. (**A**) The MEGF8 MASRPFA peptide (Figure 3B) acts as a flexible hinge in atomistic MD simulations, around which the MEGF8 ICD-MGRN1 complex can swivel. Overlay of 900 evenly distributed MASRPFA structures after alignment to the C-terminal end of the SAH. (**B**) MASRPFA Cα r.m.s.d. values are shown in light pink for each replicate, with the sliding average as a black line in. (**C**) Dictionary of Protein Secondary Structure (DSSP) matrices, show that the MEGF8 ICD and MGRN1 secondary structures remain stable during the simulations. Gray, coil; blue, β-strand; yellow, helix.

**Figure S11:**
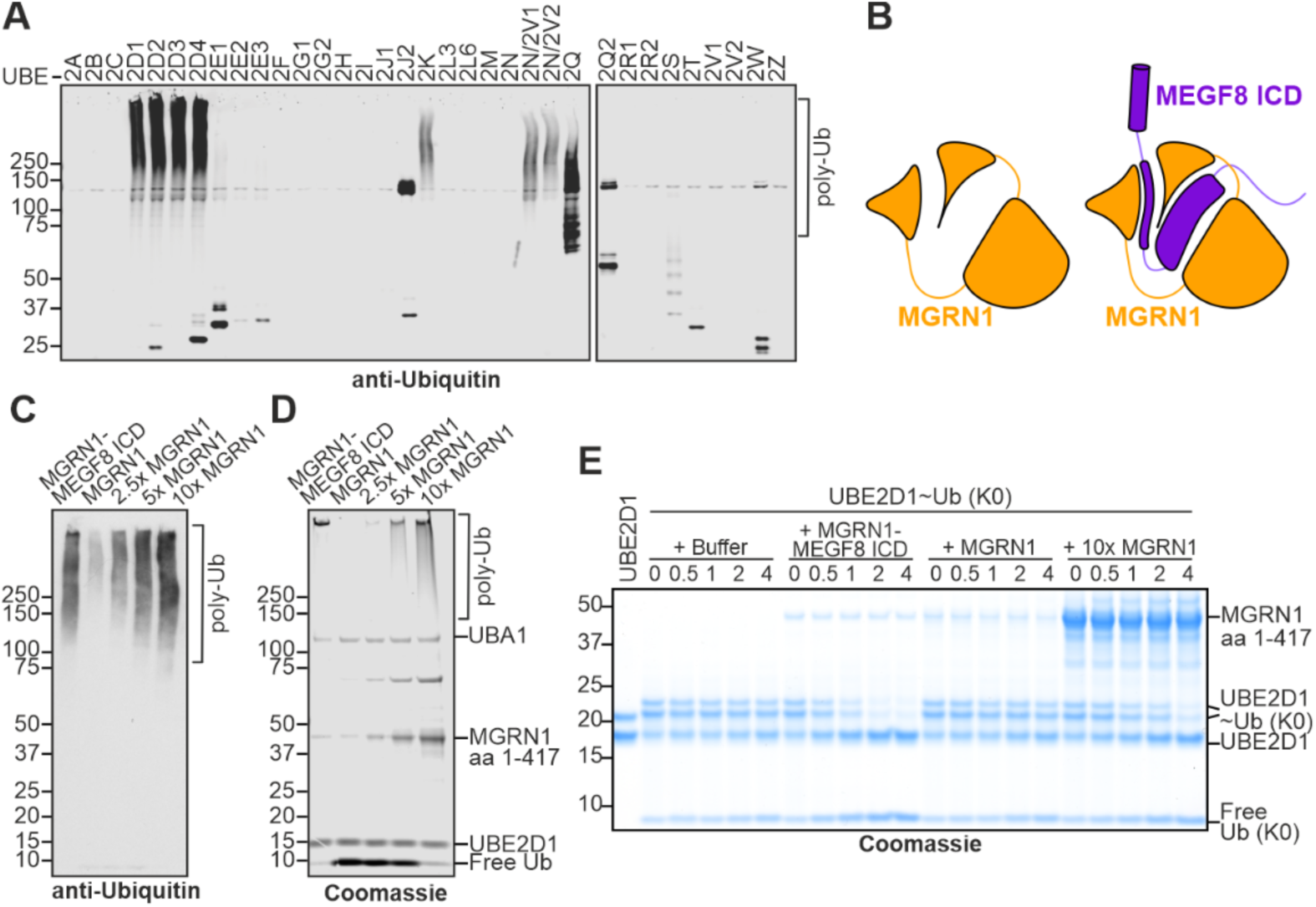
Catalytic activity of MGRN1. (**A**) A screen to identify the E2 enzyme most active in supporting poly-ubiquitylation by the MEGF8^ΔR1R2^-MOSMO-MGRN1 complex used for cryo-EM (but without the inactivating R317E linchpin mutation). E2s tested are listed across the top of the immunoblot. Reactions also contained UBA1 (E1), ubiquitin, ATP and Mg^2+^. (**B**) MGRN1 and the MEGF8 ICD-MGRN1 complex used for Ub discharge assays (and HDX-MS in Figures S9A and S9B) were both expressed in and purified from bacteria. (**C** and **D**) Poly-ubiquitin chains were detected using immunoblotting (C) or by free ubiquitin depletion (D, Coomassie stain) in reactions containing UBA1 (E1), UBE2D1 (E2), free Ub and the following E3s: MEGF8 ICD-MGRN1 complex or MGRN1 alone at the same concentration or MGRN1 alone at 2.5-fold, 5-fold or 10-fold higher concentrations. (**E**) Coomassie-stained gel showing conversion of the UBE2D1-Ub(K0) conjugate to UBE2D1 in the presence of 40 mM lysine and the indicated E3s. MGRN1 alone was used at the same concentration as the MEGF8 ICD-MGRN1 complex or at a 10-fold higher concentration. Gels like these were used to measure the kinetics of Ub(K0) discharge as shown in Figure 3K.

**Figure S12:**
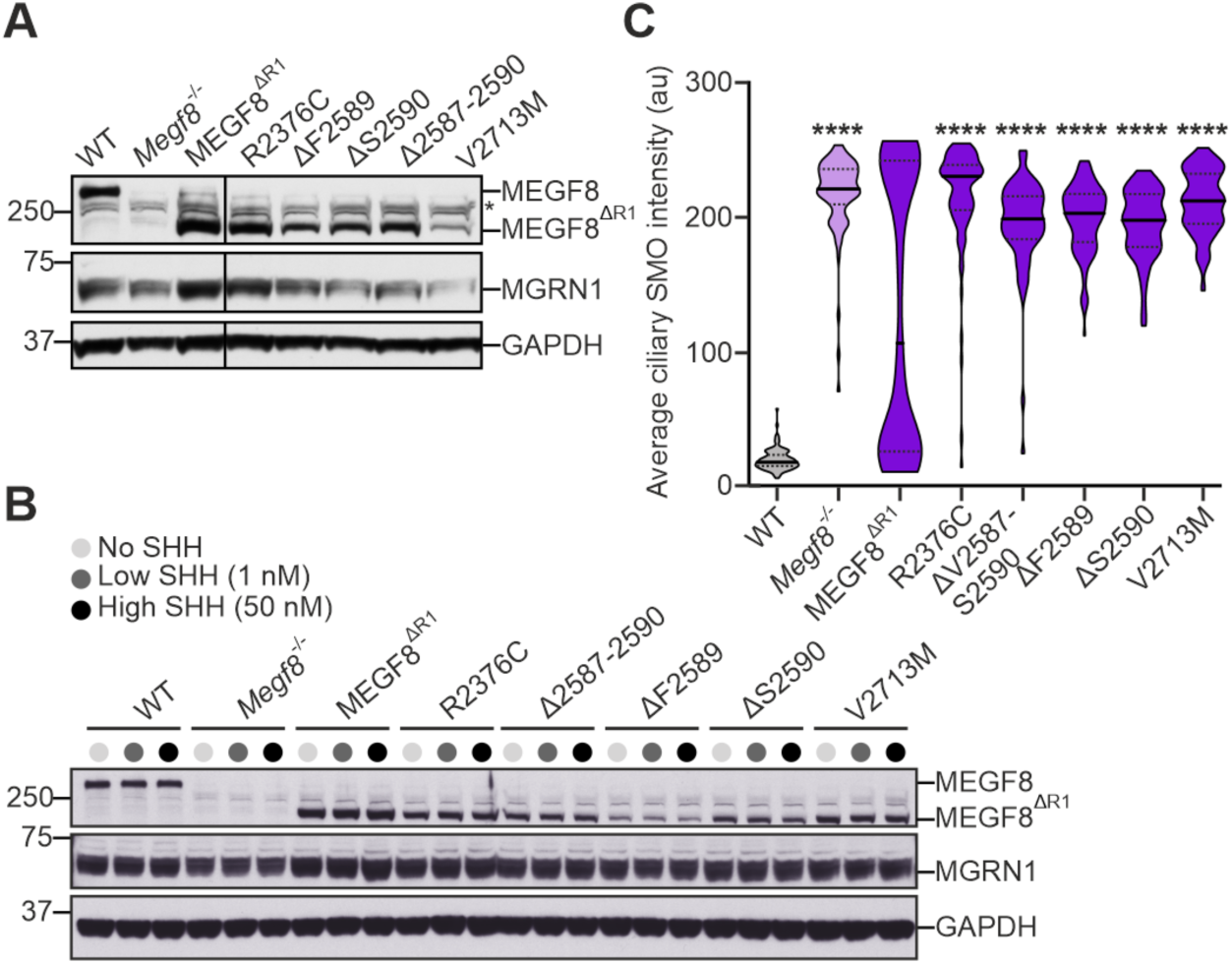
Functional analysis of Carpenter’s Syndrome mutations. (**A**) Immunoblotting was used to measure abundances of the Carpenter’s syndrome MEGF8^ΔR1^ variants functionally tested in Figures 4B and 4C. (**B** and **C**) Lentiviral transduction was used to express Carpenter’s syndrome MEGF8^ΔR1^ variants at higher levels in *Megf8*^−/−^ cells. Ciliary SMO abundance in these independently generated cell lines is shown in (C). Violin plots summarize *n*∼100 cilia and show the median (solid line) and interquartile range (dashed lines). *p*-values (****p < 0.0001) were determined using the Kruskal-Wallis (with Dunn) test in **c** (each mutant cell line compared to the WT MEGF8^ΔR1^ add back).

**Figure S13:**
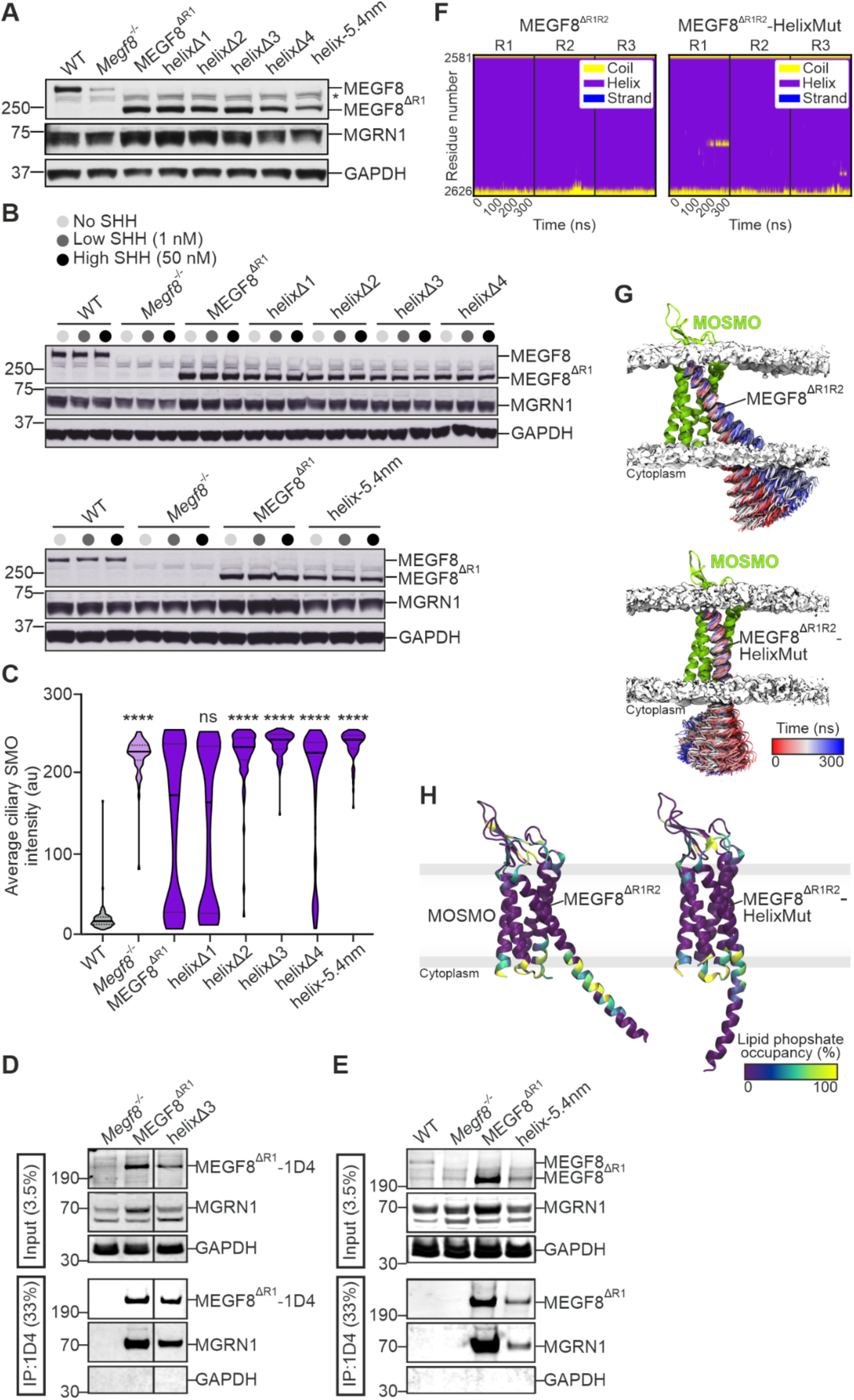
Functional and structural characterization of the central extended MEGF8 α-helix. (**A**) Immunoblotting was used to measure abundances of MEGF8^ΔR1^ SAH variants shown in Figure 5C and functionally tested in Figure 5D. (**B** and **C**) Lentiviral transduction was used to express MEGF8^ΔR1^ SAH variants at higher levels in *Megf8*^−/−^ NIH/3T3 cells. Ciliary SMO abundance in these independently generated cell lines is shown in (C). (**D** and **E**) Immunoprecipitation followed by immunoblotting was used to measure the interaction between endogenous MGRN1 with the indicated variants of MEGF8^ΔR1^, all appended with a C-terminal Rho-1D4 tag and stably expressed in *Megf8*^−/−^ cells. (**F**) DSSP matrices for three MD simulations show that the helical structure of MEGF8 SAH is maintained (despite overall conformational flexibility). (**G**) 300 evenly distributed structures of the MEGF8 central helix are colored according to simulation time. (**H**) Lipid headgroup interactions in the native (MEGF8^ΔR1ΔR2^) and ER/K stabilized (MEGF8^ΔR1ΔR2^-HelixMut) SAH structures (sequences shown in Figure S1A), quantified by per residue phospholipid occupancy calculations using the PyLipID python module^62^. MEGF8 and MOSMO are colored according to the degree of POPC lipid phosphate group occupancy around each protein residue during simulations, averaged over three replicates. Violin plots summarize *n*∼100 cilia and show the median (solid line) and interquartile range (dashed lines). *p*-values (ns>0.05, ****p < 0.0001) were determined using the Kruskal-Wallis (with Dunn) test (each mutant cell line compared to the WT MEGF8^ΔR1^ add back) (C).

**Figure S14:**
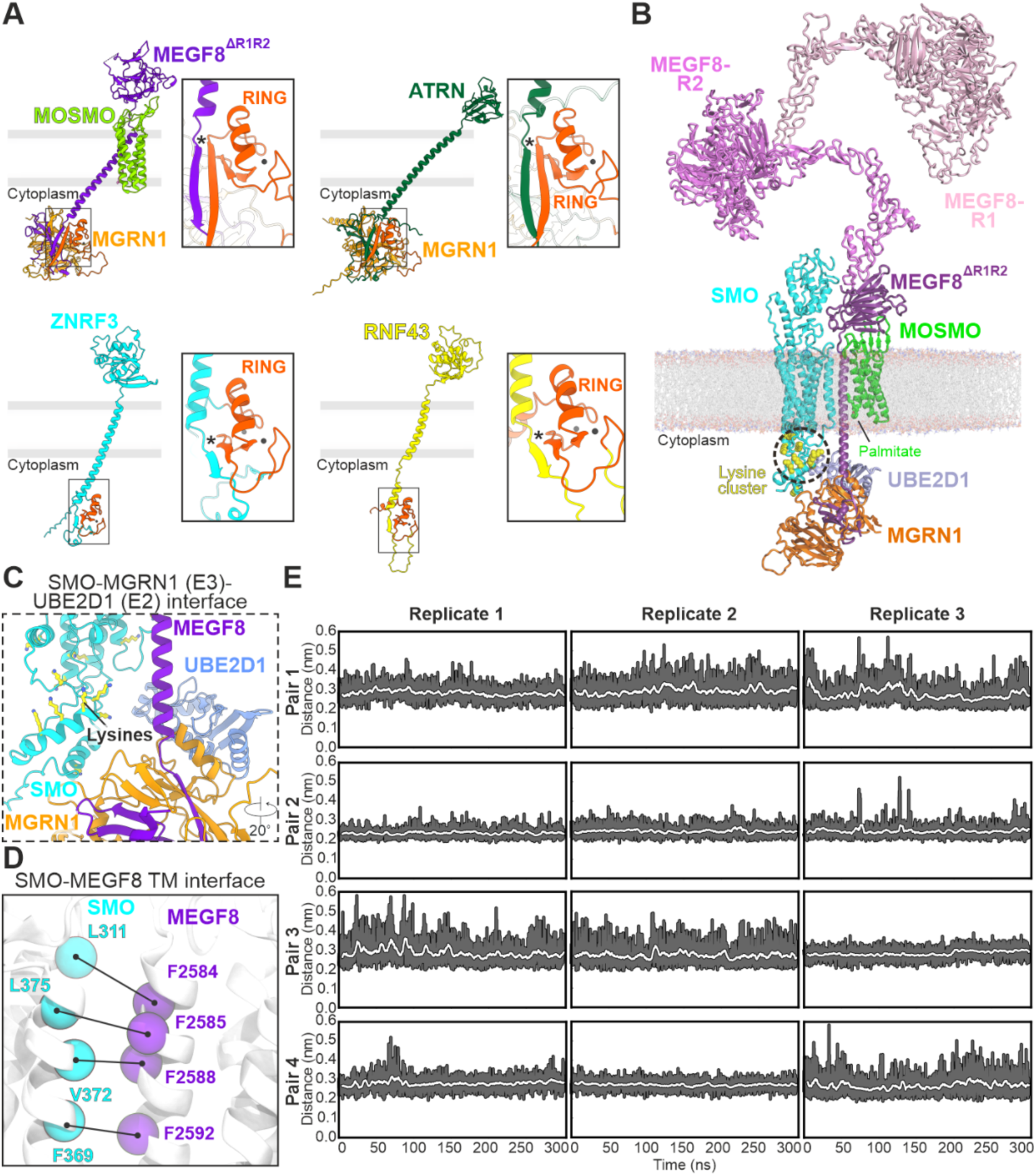
Structural model for SMO recognition by the MMM complex. (**A**) Cartoon representations of the MMM complex (this study) and AlphaFold (AF) 3 models of three related membrane-spanning E3 ligases: Attractin (ATRN)-MGRN1, ZNRF3 and RNF43. The Zn^2+^ ions in the E3 ligase RING domains (dark orange) are depicted as gray spheres. Insets show close-up views of RING domain binding and positioning by β-sheet extension (indicated by an asterisk). (**B** and **C**) Model for SMO recognition and ubiquitylation by the MMM complex. Cartoon representation of an AF2 model of the MEGF8^ΔR1ΔR2^-MOSMO-MGRN1-SMO-UBE2D1 complex, modeled in a POPC lipid bilayer using the CHARMM36 force field. AF3 models for the MEGF8 NTD (residues 1-1195 for repeat 1, R1, light purple; and residues 1143-2375 for repeat 2, R2, pink) were superimposed to the MEGF8^ΔR1ΔR2^-MOSMO-MGRN1-SMO-UBE2D1 complex. SMO lysine residues located in proximity to the MGRN1 RING domain are shown as yellow spheres (B) and using ball-and-stick representation (yellow) in the close-up view (C). These lysine residues cluster within the SMO intracellular helices 8 and 9 (B), which are unresolved in experimental structures. (**D** and **E**) The MMM-SMO interface (D) remains intact in atomistic MD simulations in a lipid bilayer. Minimum distance between SMO-MEGF8 residue pairs (P, pair 1 refers to residues SMO-311 and MEGF8-2584, pair 2 is residues SMO-375 and MEGF8-2585, pair 3 is residues SMO-372 and MEGF8-2588, pair 4 is residues SMO-369 and MEGF8-2592) as a function of time (E). Raw data for each replicate is shown in dark gray, and a sliding average is shown in white. See also **Video S3**.

## Supplementary tables

**Table S1.** Cryo-EM data collection, refinement and validation statistics. ^1,2^ Particles from two datasets collected on the same grid and microscope were merged. Total number of movies is indicated.

|  | Wild-type MMM-NB270 |  | HelixMut MMM-NB270 |  | Wild-type MM-NB270 | Wild-type MMM-NB992 |  |  |
| --- | --- | --- | --- | --- | --- | --- | --- | --- |
| Data collection |  |  |  |  |  |  |  |  |
| Microscope | Titan Krios G3i |  | Titan Krios G3i |  | Titan Krios G3i | Titan Krios G3i |  |  |
| Voltage (kV) | 300 |  | 300 |  | 300 | 300 |  |  |
| Detector | Falcon 4i – SelectrisX |  | Falcon 4i – SelectrisX |  | Falcon 4i – SelectrisX | Falcon 4i – SelectrisX |  |  |
| Recording mode | EER |  | EER |  | EER | EER |  |  |
| Magnification | 165,000 |  | 165,000 |  | 165,000 | 165,000 |  |  |
| Micrograph pixel size (Å) | 0.7303 |  | 0.7303 |  | 0.7303 | 0.7303 |  |  |
| Dose rate (e-/px/s) | 9.12 |  | 9.25 |  | 9.91 | 9.12 |  |  |
| Number of frames per movie | 60 |  | 60 |  | 60 | 60 |  |  |
| Movie exposure time (s) | 3.10 |  | 3.07 |  | 2.99 | 3.10 |  |  |
| Total dose (e-/Å)<sup>2</sup> | 50 |  | 50 |  | 50 | 50 |  |  |
| Defocus range (μM) | 1.6 to 2.6 |  | 1.6 to 2.6 |  | 1.4 to 2.6 | 1.6 to 2.6 |  |  |
| Data processing |  |  |  |  |  |  |  |  |
| No. movies/micrographs | 12,956 |  | 15,995 | 20,432<sup>1</sup> | 12,157<sup>2</sup> | 11,391 |  |  |
| Box size (px) | 480 |  | 480 |  | 384 | 480 |  |  |
| Particle number (total) | 4.2M |  | 5.1M |  | 2.2M | 1.9M |  |  |
| Particle number (final map) | 142K | 109K | 121K | 72K | 229K | 188K | 64K | 84K |
| Symmetry imposed | C1 | C1 | C1 | C1 | C1 | C1 |  |  |
| Map resolution (Å) (FSC threshold 0.143) | 2.65 | 3.33 | 2.95 | 4.41 | 2.98 | 3.29 | 3.39 | 4.66 |
| Local resolution range (Å) | 2.35 – 42.78 | 2.83 – 53.67 | 1.56 – 44.85 | 4.05 – 12.02 | 2.67 – 33.35 | 1.55 – 43.08 | 2.95 – 51.54 | 3.94 – 45.03 |
| Map sharpening B-factor (Å<sup>2</sup>) | 79.9 | 97.7 | 98.3 | 152.5 | 143.4 | 105.2 | 96.7 | 205.3 |
| Model composition |  |  |  |  |  |  |  |  |
| Protein atoms | 3923 | 7376 | 3832 | 7267 | 3854 | 7267 |  |  |
| Protein residues | 491 | 933 | 482 | 923 | 486 | 923 |  |  |
| Ligands |  |  |  |  |  |  |  |  |
| GDN sterol (AIJAX) | 1 | 1 | 1 | 1 | 1 | 1 |  |  |
| Palmitate (PLM) | 1 | 0 | 0 | 0 | 1 | 1 |  |  |
| Zn<sup>2+</sup> | 0 | 2 | 0 | 2 | 0 | 2 |  |  |
| Water | 32 | 26 | 21 | 6 | 10 | 6 |  |  |
| B-factors (Å<sup>2</sup>) – min/max/mean: |  |  |  |  |  |  |  |  |
| Protein | 19/146/62 | 0/277/42 | 25/141/68 | 25/112/150 | 16/177/79 | 22/108/150 |  |  |
| Ligand | 53/81/63 | 44/58/51 | 64/92/74 | 52/212/79 | 84/178/110 | 57/199/80 |  |  |
| Waters | 28/60/45 | 1.05/23.17/7.99 | 34/82/58 | 38/59/45 | 41/97/68 | 37/61/48 |  |  |
| RMSD from ideal: |  |  |  |  |  |  |  |  |
| Bond length (Å) | 0.003 | 0.003 | 0.003 | 0.011 | 0.003 | 0.004 |  |  |
| Bond angles (°) | 0.483 | 0.588 | 0.582 | 0.832 | 0.509 | 0.504 |  |  |
| Model validation |  |  |  |  |  |  |  |  |
| Molprobity score | 1.35 | 1.31 | 1.32 | 1.62 | 1.30 | 1.36 |  |  |
| Clashscore | 4.36 | 5.54 | 4.85 | 6.79 | 4.15 | 5.26 |  |  |
| Rotamers outliers (%) | 0 | 0 | 0 | 0 | 0 | 0 |  |  |
| FSC (0.5) model-vs-map | 2.8 | 6.4 | 3.2 | 3.4 | 3.3 | 3.4 |  |  |
| CC model-vs-map (masked) | 0.82 | 0.62 | 0.76 | 0.74 | 0.80 | 0.74 |  |  |
| Ramachandran plot |  |  |  |  |  |  |  |  |
| Favored/Allowed/Outliers (%) | 97.31/ | 98.37 | 97.67 | 96.38 | 98.15 | 97.59 |  |  |
| Allowed (%) | 2.69 | 1.63 | 2.33 | 3.62 | 1.85 | 2.41 |  |  |
| Outliers (%) | 0 | 0 | 0 | 0 | 0 | 0 |  |  |

**Table S2.** MD simulation overview. See Methods for details of model building and simulation parameters used. The protein construct without MGRN1 and the MEGF8 ICD were truncated at Ser2627 of the MASRPFA motif. This enabled a decrease in system dimensions, a concomitant decrease in computational load, and thereby access to longer timescales.

| Protein construct | Palmitoylation | Membrane (no. of POPC molecules) | System dimensions (x y z nm<sup>3</sup>) | Duration (ns) |
| --- | --- | --- | --- | --- |
| WT MEGF8<sup>ΔR1R2</sup>_MOSMO | Yes | 215 | 9 x 9 x 16 | 3 x 1000 |
| WT MEGF8<sup>ΔR1R2</sup>_MOSMO-MGRN1 | Yes | 552 | 14 x 14 x 22 | 3 x 300 |
| WT MEGF8<sup>ΔR1R2</sup>_MOSMO-MGRN1 | No | 552 | 14 x 14 x 22 | 3 x 300 |
| HelixMut MEGF8<sup>ΔR1R2</sup>_MOSMO-MGRN1 | Yes | 552 | 14 x 14 x 22 | 3 x 300 |

**Table S3.** Summary of HDX-MS experiments. Experimental conditions were optimized separately for membrane proteins (MOSMO, MEGF8) and soluble proteins (MGRN1). All experiments were carried out in duplicate, with the exception of assessing MGRN1 using membrane protein digestion protocol, as this was a control to ensure that MGRN1 did not look significantly different in the presence of detergents. Values reported here are an average of two replicates, except for the repeatability metric which is an average of the standard deviations calculated for all uptake plots shown. Values are obtained from all peptides detected across all timepoints for a given condition with medium or high confidence (based on HDexaminer fitting).

| Protein State | MGRN1 | MGRN1 (TM conditions) | MOSMO | MEGF8 |
| --- | --- | --- | --- | --- |
| # peptides | 713.5 | 454 | 46.5 | 190 |
| Sequence coverage | 99.45% | 94.00% | 60.20% | 84.60% |
| Average peptide length | 16.65 | 15.3 | 9 | 13.9 |
| Average peptide redundancy | 22.35 | 13.1 | 2 | 5.55 |
| Replicates | 2 | 1 | 2 | 2 |
| HDX reaction details: Exchange was carried out at 22 °C in 90% D<sub>2</sub>O, pH 7.4. Reaction was quenched with 2-fold dilution into urea quench solution, pH 2.4. Incubated on ice for 2 min and then flash frozen. |  |  |  |  |
| HDX time course: 15 s, 180 s, 900 s, 10800 s |  |  |  |  |
| HDX controls: Each sample was run with undeuterated control sample |  |  |  |  |
| Repeatability: Combined standard deviation for peptides assessed in uptake plots (Figures 3D-E, Figures S10F, 11C): 2.84%. |  |  |  |  |
| Back exchange: Samples were not corrected for back exchange. |  |  |  |  |

## Methods

### Cell lines

All cell lines used in this study are listed in the Key Resources Table. HEK293T Lenti-X cells were utilized to generate lentiviruses with the pHR-CMV-TetO_2_ vector^63^, which were subsequently used to infect either HEK293S GnTI^−^ TetR and Flp-In-3T3 (a derivative of NIH/3T3) cells and produce stable cell lines. HEK293T cells were also used for transient mammalian expression using the pHR-CMV-TetO_2_ vector. All adherent cell lines were cultured in Dulbecco’s Modified Eagle Medium (DMEM) containing high glucose (ThermoFisher Scientific) and supplemented with 10% fetal bovine serum (FBS) (Atlanta Biologicals), 1 mM sodium pyruvate (Gibco), 2 mM L-Glutamine (Gemini Biosciences), 1x MEM non-essential amino acids solution (Gibco), penicillin (40 U/ml) and streptomycin (40 μg/ml) (Gemini Biosciences), and cultured under standard growth conditions (37 °C, 5% CO_2_). To initiate HH signaling, NIH/3T3 cells were first ciliated by growth to confluence in DMEM containing 10% FBS followed by serum starvation in DMEM containing 0.5% FBS and the designated amount of ShhN for 24 hr. To passage cells, cells were rinsed once with PBS and then de-attached with 0.05% Trypsin/EDTA (Gemini Bio-Products). Cells were grown and maintained in standard T75 (75 cm^2^ - LentiX) or T175 (175 cm^2^ - HEK293T) flasks. HEK293S GnTI^−^ TetR cells were grown in suspension at 37 °C, 8% CO_2_, 130 rpm in FreeStyle293 medium (Invitrogen) supplemented with 1% FBS, 1% L-glutamine and 1% (v/v) NEAA with addition of no antibiotics. We used *E. coli* DH5α competent cells for cloning and *E. coli* BL21-DE3 for expression of ShhN. *E. coli* WK6 cells were used for nanobody expression. *E. coli* Rosetta2 (DE3) pLysS cells were used for expression of MGRN1 and the MGRN1-MEGF8 ICD complex.

### Constructs

Human MGRN1 (UniProt ID. O60291-4) and MOSMO (UniProt ID. Q8NHV5) genes were codon optimized for human expression and synthesized (GeneArt, ThermoFisher Scientific). MOSMO was cloned into the pHR-CMV-TetO2 vector^63^ with a C-terminal TwinStrep tag^64^ or Rho1D4 antibody epitope tag^65^. MGRN1 was cloned into the pHR-CMV-TetO2 vector resulting in a C-terminal 6xHis tag. Human MEGF8 (UniProt ID: Q7Z7M0-2)^10^ served as a template for generating all MEGF8 variants, which were subsequently cloned into the pHR-CMV-TetO2 vector in frame to either a C-terminal Rho1D4 tag or a 3C protease cleavage site followed by a TwinStrep tag. MEGF8^ΔR1R2^-HelixMut was created using an stabilized α-helical motif^29,66^. All point mutants of MEGF8, MOSMO or MGRN1 were generated by Gibson Assembly and subsequent Gateway Cloning into a shuttle vector (Gateway™ pENTR™ 2B, ThermoFisher Scientific) or Gibson Assembly within their respective expression vectors. MGRN1 constructs were cloned into pETDuet-1 vector for dual expression in bacteria using the backbone from Addgene #133322^67^. Constructs for expressing UBE2D1 and UBA1 in bacteria were generously gifted by the Hay laboratory^26,68^. The ubiquitin (Ub, K0) construct for bacterial expression, a generous gift from Rachel Klevit, encodes a lysine-free ubiquitin with the following mutations: Lys6Arg, Lys11Arg, Lys29Arg, Lys33Arg, Lys48Arg, Lys63Arg and Lys27Met^27^.

### Cell line generation and expression of membrane proteins

Stable cell lines for production of membrane proteins were established by following a previously published protocol^63^. MEGF8-MOSMO and MEGF8-MOSMO-MGRN1 (MMM) complexes were formed by co-expression. Stably-transfected cells were grown to densities of 2-3 ×10^6^ cells/mL and induced with doxycycline (Merck). Media was harvested by centrifugation (1,500 xg, 4 °C, 15 min) after 72 hours, except for MOSMO, which was harvested at 48 hours. Pellets were flash-frozen and stored at −80 °C. For small-scale screening, HEK293T cells in 6-well plates were transiently transfected using polyethylenimine (PEI) and harvested after 48 hours.

### Purification of membrane proteins

All steps were performed at 4 °C. For purification of MOSMO and MEGF8-MOSMO complexes, frozen cell pellets were thawed and resuspended in HEPES buffer. Membranes were solubilized and cell debris was removed by centrifugation (40,000 xg, 30 min, 4 °C). Supernatant was diluted 2-fold in lysis buffer and purified via Rho-1D4 antibody affinity. MEGF8^ΔR1R2^-MOSMO complex samples for hydrogen/deuterium exchange mass spectrometry (HDX-MS) were purified in a similar manner to above. For the MMM complex, tandem purification was applied using StrepTactinXT 4Flowaffinity followed by TALON bead purification. For all samples, the eluates from the affinity columns were concentrated and further purified via size-exclusion chromatography (SEC) using a Superose 6 Increase 10/300 GL column.

### Protein expression in bacteria

MGRN1 alone and the MEGF8 ICD -MGRN1 complex were expressed in Rosetta™ 2 (DE3) pLysS Competent cells (Millipore-Sigma) grown to an optical density of 0.8-1.0 in terrific broth supplemented with kanamycin or carbenicillin at 37 °C, at 195 rpm. Cultures were induced with 0.5 mM isopropyl β-d-1-thiogalactopyranoside (IPTG) and proteins were expressed overnight at 18 °C. Cells were harvested by centrifugation (4,000 xg, 15 min) and resuspended in lysis buffer Pellets were flash-frozen and stored at −80 °C. For expression of MGRN1 alone, the protein was expressed with N-terminal 6xHis-SUMO tag. For expression of the MEGF8 ICD-MGRN1 complex, MGRN1 was expressed with an N-terminal MBP-SUMO tag and the MEGF8 ICD with an N-terminal 6xHis tag.

### Protein purification from bacteria

All steps were carried out on ice or at 4 °C. Bacterial pellets were thawed in a lysis buffer, lysed by homogenization (EmulsiFlex-C5, Avestin), and clarified by centrifugation. Supernatants were collected and incubated with cOmplete^™^ His-Tag Purification Resin (Millipore-Sigma) for 2 hours. The resin was collected on a gravity flow column and washed extensively with Tris-HCl buffer. For His-SUMO tagged MGRN1 alone, beads were incubated overnight with SENP1 protease (0.5 mg/L of bacterial culture). MGRN1 was purified from the flow-through using a RESOURCE Q anion exchange column (Cytiva). For pellets expressing both MBP-SUMO-MGRN1 and 6xHis-MEGF8 ICD, we used sequential purification on His-Tag purification resin followed by Amylose resin to obtain the complex. Proteins were eluted from the His resin in 20 mM Tris pH 8, 500 mM NaCl, 1 mM TCEP, 250 mM imidazole, 5% (v/v) glycerol. The eluate was then incubated with Amylose resin (NEB) to capture the complex. To cleave the SUMO tag from MGRN1 protein, the sample was incubated with SENP1 protease ^69^ and the flow-through was collected. As a final polishing step, all proteins (MGRN1 alone and the MEGF8 ICD-MGRN1 complex) were fractionated on a HiLoad 16/60, Superdex 200 prep grade or a Superose 6 10/300 GL (Cytiva) column. Fractions from a monodisperse peak (and separated from the void volume) were pooled, concentrated, flash-frozen, and stored at −80 °C. UBE2D1, UBA1, and Ub (K0) were expressed and purified as described previously^26,68,70^.

### Generation of nanobodies

For nanobody immunization, the MEGF8^ΔR1R2ICD^-MOSMO complex was purified as described for the MEGF8^ΔR1R2^-MOSMO complex. Peak fractions were pooled, concentrated to 1 mg/mL, and flash-frozen.

The immunization and Nanobody library construction were carried out according to a previously published protocol ^71^. cDNA libraries from Llama peripheral blood lymphocytes were cloned into golden gate variant of the pMESy4 phagemid vector (GenBank: KF415192) containing an N-terminal pelB signal peptide and a C-terminal 6xHis and EPEA tags. Nanobodies against the MEGF8^ΔR1R2ICD^-MOMSO complex were selected by phage display using established protocols^71^.

Nanobodies were expressed and purified as described previously^72^.

### Cryo-EM sample preparation

The MMM and MEGF8-MOMSO complexes were incubated with the purified nanobody at 4 °C. Complexes were concentrated and purified by SEC. Samples were concentrated and subsequently used for cryo-EM grid preparation. Prior to data collection, 3.5 μL sample was applied to a 300-copper mesh Quantifoil Holey Carbon grid R 1.2/1.3 without glow-discharging. All grids were prepared using a Vitrobot mark IV (FEI) at 4 °C and 100% relative humidity.

### Cryo-EM data collection

Data were collected at the Oxford Particle Imaging Centre (OPIC) on a 300 kV Titan Krios G3i microscope with Falcon4i direct electron detector and SelectrisX energy filter (ThermoFisher Scientific) and at electron Bio-Imaging Centre (eBIC). Automated data collection was set up in EPU (ThermoFisher Scientific). Multi-grid EPU (ThermoFisher Scientific) was used to collect data of the MEGF8^ΔR1R2ICD^-MOMSO complex with four different nanobodies, to select suitable nanobodies (NB270 and NB992) for high-resolution structure determination. Data collection parameters for different samples are outlined in **Table S1**.

### Cryo-EM image processing

All datasets were processed using cryoSPARC v.4.2.1-4.6.2 ^53^ following the workflows shown in **Figure S2.** For each dataset, raw movies were aligned by Patch Motion Correction. The contrast transfer function (CTF) was estimated using Patch CTF and low-quality micrographs were subsequently discarded after manual inspection (e.g., astigmatism, ice contamination). Particles were initially blob picked, and junk particles were removed manually. The binned particles with a pixel size of ca. 2.739 Å/px were subject to 2D classification, where the best 2D class averages showing different views and secondary structure elements were selected for template picking. After generating ab-initio models, the particles from the best-resolved class were selected for Topaz training using the full dataset^73^. All particles belonging to classes displaying MEGF8-MOSMO or MMM complexes were merged, duplicate particles removed, and re-classified by heterogenous refinement using three or more classes. The particles from the best class were re-extracted in a 480 px box size (0.7303 Å/px), before running non-uniform refinement. To separate low resolution MEGF8-MOSMO particles, 3D classification without alignment was performed using a soft mask surrounding the extracellular and transmembrane regions (including NB270 or NB992). Per-particle defocus was refined for particles belonging to the highest resolution class, before locally refining with the same mask. To identify particles with MGRN1, a soft mask surrounding only MEGF8 ICD and MGRN1 protein residues was created by simulating density from an AlphaFold (AF) model using the ChimeraX molmap function. This mask was used for 3D classification without alignment, then particles belonging to the best class were locally refined with a mask encompassing MGRN1 and the transmembrane domain (TMD). Local resolution was estimated in cryoSPARC and rendered in ChimeraX-1.9. The local refinement maps were combined in PHENIX to create a composite map^55^. See **Supplementary note 1** for further details.

### Model building, refinement and validation

Initial models for the MMM complex were generated using AF2^74^, implemented in ColabFold^75^, and reanalyzed with AF3 through the DeepMind portal (https://alphafoldserver.com)^28^. An AF3 model of the MMM complex and an AF2^74^ model for NB270 were placed into the wild-type MMM-NB270 map using the ChimeraX-1.4 ‘fit-in-map’ function. One cycle of rigid body real space refinement in PHENIX was performed to optimize the overall fit of the model to the map. Model geometry was improved by multiple cycles of real-space refinement in PHENIX, interspersed with manual rebuilding in Coot^76^. The GRADE server was used to generate a restraint dictionary for the GDN molecule and palmitoyl. Distance between MOSMO Cys164 SG and palmitoyl C1 atoms was restrained to 1.77 Å using custom restraints. This model was then used to refine the MMM-NB270 HelixMut, MEGF8^ΔR1R2ICD^-MOSMO-NB270 and MMM-NB992 complexes in a similar manner. For all models, model geometry was validated using MolProbity^77^. Map and model statistics are listed in **table S1**. Sequence conservation was calculated using Consurf^78^. Buried surface areas of protein–protein interfaces were calculated using the PISA webserver^34^ and shape complementarity statistic Sc in CCP4^32^. Data analysis and visualization were conducted using ChimeraX version 1.4 to 1.9^79^ and PyMOL (Schrödinger, LLC.).

### AlphaFold modeling

For MD simulations in **Figure S14**, an AF2^74^ model of the 1:1:1:1:1:1 human SMO (UniProt ID: Q99835, residues 60-652): MEGF8 (residues 2374-2749): MOSMO: MGRN1 (residues 18-377): UBE2D1 (UniProt ID: P51668): Ub complex was generated using the Tamarind Bio utility (https://app.tamarind.bio/tools/alphafold). The extracellular domain (ECD) of MEGF8 was modeled using AF3 and superimposed to the AF2 model of the MMM-Smoothened (SMO)-E2 complex (**Figure S14B**). The AF3^28^ models for MEGF8-repeat 1 and MEGF8-repeat 2 were merged in Coot. The schematic in **Figure 5H** was guided by AF3 models of full-length ZNRF3, RNF43, MARCH6 and the Attractin (ATRN)-MGRN1 complex.

### Atomic molecular dynamics simulations

All simulations were performed using GROMACS v2023.3 and v2024.3^80^. The unresolved C-terminal residues Leu165, Pro166, and Gly167 of MOSMO were added using PyMOL (Schrödinger, LLC). Cys46-Cys56 within MOSMO, and Cys2394-Cys2385 within the Stem domain of MEGF8 were treated in their disulfide linked forms. Cysteines forming the Zn^2+^ ion binding sites of MGRN1 were treated in their thiolate forms. The side chain of Asp16 in MOSMO was treated as protonated due to its exposure to the hydrophobic core of the bilayer^81^. Models were embedded into 1-palmitoyl-2-oleoyl-sn-glycero-3-phosphocholine (POPC) lipid bilayers using CHARMM-GUI membrane builder^82^. Systems were solvated with TIP3P waters and neutralized using 0.15 M NaCl. Energy minimization was performed using the steepest descents algorithm implemented in GROMACS. Equilibration was achieved with harmonic position restraints applied to Cα atoms and lipid atoms. Restraints were faded out over six steps following the CHARMM-GUI equilibration procedure^83^. The final step employed only position restraints on the Cα atoms, and the equilibration duration was modified to 10 ns. Energy minimization and equilibration procedures were performed independently for each repeat using different random initial velocity seeds. Systems were subsequently subjected to between 300 and 1000 ns of unrestrained atomic molecular dynamics (**Table S2**).

Simulations were run using the CHARMM36m^84^ and CHARMM36^85^ force fields. Temperature was controlled at 303.15 K using a V-rescale thermostat^86^ (τt = 1.0). Pressure was maintained at 1 bar using a C-rescale barostat^88^ (τp = 5.0) with semiisotropic coupling and a compressibility of 4.5 × 10-5 bar^−1^. Periodic boundary conditions were applied. Long-range electrostatics were described using Particle Mesh Ewald (PME) ^87^ with a 12 Å cutoff. Van der Waals interactions were calculated using the Verlet method with a 12 Å cutoff. Equations of motion were integrated using a 2 fs time step. The LINCS algorithm was used to constrain covalent bond lengths^88^. Simulations were analyzed using tools available in GROMACS^80^, Visual Molecular Dynamics (VMD)^89^, PyMOL (Schrödinger, LLC), PyLipID^62^, and MDTraj^90^.

### Stable cell line generation for signaling assays

Stable expression of tagged MMM components in their respective knockout NIH/3T3 line was carried out by either lentiviral expression or Flp-mediated DNA recombination into a single defined locus in the genome (Flp-In system, ThermoFisher Scientific), as described previously^10,37^.

### Western blotting

Cell lines were seeded in 6 cm tissue culture plates at 550,000 cells per dish in high glucose, complete DMEM. After 48 hours, the FBS concentration was reduced to 0.5% (v/v) and cells were treated with either no Sonic Hedgehog (SHH), 1 nM SHH or 50 nM SHH for 24 hours. SHH was produced as described previously^91^. Cells were harvested in RIPA lysis buffer (50 mM Tris-HCl pH 7.4, 150 mM NaCl, 2% (v/v) NP-40, 0.25% (v/v) Deoxycholate, 0.1% (w/v) SDS, 1mM DTT, 1 mM NaF, 1 mM Na_3_VO_4_, 10% (v/v) glycerol, 1x SIGMAFAST protease inhibitor cocktail (Sigma-Aldrich)), before assessing protein concentration using Pierce™ BCA Protein Assay Kit (ThermoFisher Scientific). Samples (containing 20-40 µg of total protein) were prepared in 1x NuPAGE LDS Sample Buffer (ThermoFisher Scientific) and 50 mM DTT, and then incubated at 37 °C for 30 min. Samples were loaded onto a NuPAGE™ Bis-Tris 4-12% gel (ThermoFisher Scientific) and run at 125 V for 2 to 3 hours in NuPAGE™ MOPS SDS Running Buffer (ThermoFisher Scientific). Proteins were transferred to nitrocellulose membranes (Bio-Rad), and then blocked in 5% (w/v) milk in Phosphate Buffered Saline with 0.1% (v/v) Tween and incubated with primary antibody overnight at 4 °C. The following day, membranes were incubated with secondary antibody for 1 hour at room temperature. Membranes were visualized using Pierce™ ECL Western Blotting Substrate (ThermoFisher Scientific) and exposed to film for imaging.

### Antibody information

Commercial primary antibodies included Ub (P4D1, Santa Cruz Biotechnology, sc-8017, 1:1000), His-tag Antibody (Cell Signaling Technologies, #2365, 1:1000), RNF156 (MGRN1) Polyclonal Antibody (Proteintech, #11285-1-AP, 1:2000), GAPDH (Proteintech, 60004-1-Ig, 1:100,000), HA Tag monoclonal antibody (Proteintech, 66006-2-Ig, 1:3000), Rho 1D4 Antibody (University of British Columbia), FLAG tag Antibody (Sigma Aldrich, MAB3118, 1:3000), Anti-p38 Antibody (Abcam, ab7952), and FK2 antibody against mono- and poly-ubiquitylated conjugates (Ubiquigent, #68-0121-500). For PLA, Mouse Anti-FLAG (Invitrogen, MA1-91878), and Mouse Anti-3xHA (Invitrogen PI26183) were used. The following primary antibodies were generated in the lab: Guinea pig anti-ARL13B (1:1000)^92^; rabbit anti-SMO (designed against an intracellular epitope, 1:2000)^93^; rabbit anti-MEGF8 (1:250)^10^. Secondary antibodies included Peroxidase AffiniPure® Donkey Anti-Rabbit or Anti-Mouse IgG (H+L) (Jackson ImmunoResearch) (for Western blot), or Donkey Anti-Guinea Pig or Rabbit IgG (H+L) coordinated to AlexaFluor dyes (ThermoFisher Scientific).

### Immunofluorescence

Ciliary SMO was analyzed by immunofluorescence as described^12^ with the exception that samples were incubated with primary antibody overnight. Images were collected^12^ and analyzed with CiliaQ^94^. During the CiliaQ preparator phase, images were background subtracted with a radius of 10 and smoothed with a radius of 0.5. RenyiEntropy method was used for segmenting the Cilia channel. Images were then manually inspected to ensure that the segmented channel overlapped with identifiable cilia, based on Arl13b staining. Any manual corrections were made using the CiliaQ editor. SMO intensity was measured for each cilia identified in the segmented channel, removing any cilia less than 100 voxels or touching the x or y border of the image. An increased range function for connecting cilia intensity regions was allowed by 1 voxel. Voxels that are detected in the cilia channel that are diagonally adjacent to a detected cilia object will be counted as part of the cilia region.

### *In situ* Proximity Ligation Assay (PLA)

Cells were cultured on sterile coverslips at 80,000 cells per well in complete DMEM for 48 hours, then switched to fresh DMEM containing 0.5% (v/v) FBS. After 24 hours, cells were washed with PBS and fixed with 4% (v/v) PFA in PBS for 10 min. PLA was performed using the Duolink® *In Situ* Orange Starter Kit Mouse/Rabbit kit (DUO92102, Millipore-Sigma) following the manufacturer’s instructions. Briefly, cells were blocked with blocking solution for 1 hour at 37 °C and incubated with primary antibodies for 1 hour at 37 °C. All primary antibodies were diluted to 1:100 in blocking buffer prior to use. Cells were then incubated with the Duolink® *In Situ* PLA® Anti-Rabbit PLUS and Duolink® *In Situ* PLA® Anti-Mouse MINUS probes for 1 hour at 37 °C. Finally, ligation and amplification were performed for 30 min and 100 min, respectively, before mounting the cells in ProLong Diamond Antifade Mountant with DAPI (P36962, Invitrogen).

### Imaging, quantification, and statistical analysis of PLA images

Fluorescent images were acquired on the Nikon Eclipse Ti2 Microscope equipped with a 60x oil immersion objective (NA 1.42). Z-stacks (0.3 µM sections) were acquired with identical gain, offset, and laser power settings used for all conditions within an experiment. The saved ND2 files were then opened in Fiji 1.54f (ImageJ2) with Z-stack projections of maximum fluorescent intensities. To determine the mean PLA signal in each cell, the number of PLA foci were adjusted with the number of cells to obtain the relative PLA dots/cell value. GraphPad Prism 8 was used to analyze and plot the data.

### Real Time Quantitative Reverse Transcription PCR (qRT-PCR)

RNA was extracted from 3T3 cells using TRIzol reagent (Life Technologies), followed by cDNA synthesis using iScript Reverse Transcription Supermix (BioRad). qRT-PCR was performed on QuantStudio 5 Real-Time PCR System (ThermoFisher Scientific) with custom primers *Gli1* (Fwd: 5’-ccaagccaactttatgtcaggg-3’ and Rev: 5’-agcccgcttctttgttaatttga-3’), and *Gapdh* (Fwd: 5’-agtggcaaagtggagatt-3’ and Rev: 5’-gtggagtcatactggaaca-3’). ΔΔCt was determined relative to *Gapdh* transcript levels.

### *In vitro* auto-ubiquitylation assays

Reactions were prepared in 50 mM HEPES pH 7.4, 150 mM NaCl, 0.5 mM TCEP, 5% (v/v) glycerol with 250 nM E3 or E3 complex, 20 µM Ub (R&D Systems, U-100H), 2 µM UBE2D1, and 100 nM UBA1 (final concentrations in reactions)^26,68^. To begin the reaction, ATP and MgCl_2_ were added to a final concentration of 2 mM and 5 mM, respectively. Reactions were quenched with 1x LDS Sample Buffer (ThermoFisher Scientific), 33 mM DTT and 25 mM EDTA. Samples were then loaded onto a NuPAGE Bis-Tris 4-12% gel for subsequent analysis. Gels were analyzed by Western blot, as described above, or by Coomassie staining. E2 scan was performed using E2 Scan Kit version 2 (Ubiquigent, Cosmo Bio). All reagents for the auto-ubiquitylation reaction were provided within the kit except for the E3, which we generated as described above.

### *In-vitro* Ub discharge assays

UBE2D1∼Ub(K0) conjugates were prepared and purified by SEC, as described previously^70^. The UBE2D1∼Ub(K0) conjugates (13 µM) were mixed with either the MEGF8 ICD-MGRN1 complex (0.7 µM) or MGRN1 at 0.7 µM (same concentration as the complex) or 7 µM (10-fold higher concentration than the complex). Note that concentrations are based on equalizing the intensity of the Coomassie Blue stained band for MGRN1 in both E3 preparations. At time = 0, reactions were started by adding L-Lysine (40 mM) and samples were moved to 37 °C. Reactions at each time point were stopped by adding 1x LDS Sample Buffer (ThermoFisher Scientific) and subsequently analyzed by SDS-PAGE.

### Hydrogen-deuterium exchange labeling reaction

Proteins were prepared at a stock concentration of 10 μM in their respective SEC buffer. D_2_O buffer was prepared by lyophilizing SEC buffer and resuspending in D_2_O (Sigma Aldrich). For protein samples containing GDN, D_2_O buffer was prepared without detergent to limit the amount of detergent in the final sample. To begin the labeling reaction, samples were diluted to 1 μM in D_2_O buffer. The labeling reaction was quenched by mixing samples 1:1 with Quench buffer (3 M Urea, 20 mM TCEP, pH 2.4). The samples were mixed, returned to ice for 2 min and then flash frozen in liquid N_2_.

### Liquid chromatography/mass spectrometry (LC/MS) analysis

LC/MS methods, peptide identification and analysis were performed as described previously^95^. Briefly, samples were thawed and immediately injected on a cooled valve system (Trajan LEAP) coupled to liquid chromatography (Thermo Ultimate 300) flowing Buffer A (0.1% formic acid). After injection, samples were flowed onto protease columns (Nepenthesin-2-pepsin (Affipro) for transmembrane samples, or porcine pepsin and aspergillopepsin columns prepared the lab for soluble proteins^96^), followed by desalting onto a manually packed trap column, over 3 min. Following desalting, peptides were separated over a C18 analytical column by acetonitrile gradient increasing 5% to 45% over 17 min, followed by 45%-90% over 1 min. After separation by liquid chromatography, peptides were eluted directly onto a Q Exactive Orbitrap Mass Spectrometer (ThermoFisher Scientific) operating in positive ion mode (MS1 settings: resolution 140000, AGC target 3e6, maximum IT 200 ms, scan range 300-1500 m/z). Initially, peptides were identified for each protein by running MS/MS on an undeuterated sample (MS1 same as above, MS2 settings: resolution 17500, AGC target 2e5, maximum IT 100 ms, loop count 10, isolation window 2.0 m/z, NCE 28, charge state 1 and >8 excluded, dynamic exclusion 15.0 s). For transmembrane proteins, a blank injection with shorter gradient was performed between every sample.

Analysis of resulting peptides was carried out as described previously^95^. Briefly, a list of reference peptides was determined based on MS/MS data processing in Byonic (Protein Metrics). HD-Examiner (Version 3.1, Sierra Analytics) was used to analyze deuterium uptake data based on these reference peptides. Settings were adjusted to account for 1:10 dilution of the undeuterated sample into a deuterated buffer, but otherwise default settings were utilized. For unimodal peaks, the difference in mass centroid between undeuterated and deuterated peaks were used to determine changes in deuterium uptake. For bimodal peaks, the spectra were fit (as described previously^96^) globally to sums of Gaussians such that the widths were constant across time points. Fitting procedure was adapted from a previous study^96^. See **Table S3** for further details.

### Intact mass spectrometry analysis

To identify MOSMO palmitoylation, intact mass spectrometry of detergent solubilized wild-type and C164S MOSMO were performed as described previously^97^. Wild-type and MOSMO-C164S were purified as described above using StrepTactinXT 4 Flow resin. MOSMO was then passed over a Superose 6 Increase 10/300 GL column (Cytiva) equilibrated in 20 mM HEPES pH 7.5, 150 mM NaCl, 0.03% (w/v) DDM, and 0.006% (w/v) CHS. Peak fractions were pooled, concentrated to 1 mg/mL, and flash-frozen prior to intact mass spectrometry.

### Statistical analysis

Statistical significance for differences in ciliary SMO abundance, measured by immunofluorescence and quantified using CiliaQ (see above), was determined using the non-parametric Kruskal-Wallis test, followed by Dunn’s multiple comparison test. For each cell line >100 cilia were analyzed from >5 independent images and the entire experiment repeated using a different batch of cells (with similar results). Quantitative RT-PCR was performed three independent times, each with three technical replicates and significance of any differences assessed by one-way ANOVA with Dunnett’s multiple comparison test. To assess the statistical differences in PLA data, we used one-way ANOVA. Note that PLA data passed the Kolmogorov-Smirnov test for normality.

## Supplementary notes

### Note 1: Detailed cryo-EM structure solution of MM and MMM complexes

All datasets were processed using cryoSPARC^53^. For the wild-type MMM-NB270 complex, a total of 4.2 M particles were picked from 12,956 micrographs and extracted in a 480 px box Fourier cropped to 128 px (2.739 Å/px) (**Figure S2A**). After template picking, 4 ab-initio models were generated and refined by heterogenous refinement. This process yielded 2 main classes of the MMM complex, with 1:1 or 1:2 MEGF8 to MOSMO stoichiometries, 1 class of the binary MEGF8-MOSMO complex, and 1 junk class. The particles from each model were used to train the neural network in Topaz^73^. After one round of Topaz picking, duplicate particles were removed, yielding 512,688 particles. The particles were re-extracted in a 480 px box, before running non-uniform refinement. For the MEGF8-MOSMO subcomplex, a mask was created around extracellular and transmembrane regions of MEGF8-MOSMO-NB270. 3D classification without alignment was then performed on the unbinned particles with 4 classes, leading to a best-resolved class with 142,461 particles. Non-uniform refinement of this class resulted in a map resolution of 2.74 Å and subsequently, these particles were CTF refined by local CTF refinement. Following subtraction of the detergent micelle, particles were locally refined to 2.65 Å. For the MMM complex, multiple rounds of Topaz were performed to find more particles with additional density for MGRN1. A total of 554,435 unbinned particles were subject to 3D classification with 5 classes and a mask encompassing MGRN1. Non-uniform refinement of the best-resolved MGRN1 class with 109,449 particles resulted in a map resolution of 3.52 Å. These particles were subsequently processed by local refinement to mask out the detergent micelle, resulting in an overall map resolution of 3.33 Å. The local refinement maps were combined in PHENIX to create a composite map^55^.

For the wild-type MMM-NB992 complex, a total of 1.9 M particles were picked from 11,391 micrographs and extracted in a 480 px box Fourier cropped to 128 px (2.739 Å/px) (**Figure S2B**). Particles were initially blob picked in cryoSPARC Live, before selecting 310,076 particles from one round of 2D classification. 3 ab-initio models were generated and refined by heterogeneous refinement, yielding a junk class and two major classes of the MMM complex with 1:1 or 1:2 MEGF8 to MOSMO stoichiometries. Blob picking was also performed on all motion-corrected micrographs in cryoSPARC and 563,550 particles were selected from 2D classification. One round of ab-initio coupled to heterogeneous refinement was performed on these selected particles. Those particles with a 3D class posterior probability lower than 90% in the best-resolved class (97,983 particles) were discarded. The resulting 130,762 particles, with a 1:1 MEGF8 to MOSMO stoichiometry in the MMM complex, were used to train Topaz picking. After one round of Topaz picking, 551,368 particles were selected by 2D classification. 3 ab-initio models were generated and refined by heterogeneous refinement. Duplicate particles from all of the best-resolved 3D classes were removed, yielding 345,808 particles. The particles were re-extracted in a 480 px box Fourier cropped to 300 px (1.168 Å/px), before further classifying particles into 2 volumes using ab-initio reconstruction. Those particles belonging to the highest resolution class were unbinned and non-uniform refinement resulted in a map with a resolution of 3.58 Å. Particles were locally refined using a mask around the extracellular regions of MEGF8-MOSMO and NB992, resulting in a focused refinement map with an overall resolution of 3.29 Å. For the MEGF8-MOSMO subcomplex, a mask was created around extracellular and transmembrane (TM) regions of MEGF8-MOSMO-NB992. 3D classification without alignment was then performed on the unbinned particles with 3 classes, leading to a best-resolved class with 64,250 particles. These particles were locally refined to 3.39 Å. For the MMM complex, particles were subject to 3D classification with 3 classes and a mask encompassing MGRN1. After particle subtraction of the detergent micelle, particles were locally refined using a mask encompassing the TM and cytoplasmic regions of the map, resulting in an overall map resolution of 4.66 Å. These three local refinement maps were combined in PHENIX to create a composite map.

For the MMM-NB270 HelixMut complex, 3.4 M particles were extracted in a 480 px box Fourier cropped to 128 px (2.739 Å/px) from 15,995 micrographs (**Figure S2C**). After 3 rounds of 2D classification, 5 ab-initio models were generated from the binned particles and re-classified by heterogeneous refinement. The major class contained 327,128 particles with 1:1 MEGF8 to MOSMO stoichiometry, which were subsequently re-extracted in a 480 px box (1.168 Å/px). Non-uniform refinement on this set of particles resulted in a map with a 3.75 Å resolution. For the MEGF8-MOSMO complex, 3D classification without alignment was performed using a mask encompassing NB270 and the extracellular and TM domains of MEGF8 and MOSMO. This sorted particles into 3 classes, yielding a major class with 120,692 particles. These particles were then re-extracted in a 480 px box (0.7303 Å/px) and refined by non-uniform refinement, resulting in a map with 3.07 Å overall resolution. Following particle subtraction of the detergent micelle and cytoplasmic regions (MEGF8 ICD and MGRN1) of the map, particles were locally refined to 2.95 Å using the same mask as for 3D classification without alignment. For the MMM complex, an additional 4,437 movies were collected on the same grid and processed in the same manner. 228,675 particles were extracted in a 480 px box Fourier cropped to 128 px (2.739 Å/px), increasing the number particles in the heterogenous refinement map (327,128 total images in the dataset) to 615,803 particles. The particles from both datasets were classified together into 4 classes by heterogenous refinement, before one round of Topaz picking. Subsequently, the unduplicated particles were classified by a second round of heterogeneous refinement. The best-resolved class for MGRN1 contained 211,378 particles, which were re-extracted in a 480 px box Fourier cropped to 300 px (1.168 Å/px). This set of particles was processed by 3D classification into 3 classes, without alignment. 3D classification was run using a mask around MGRN1. Following 3D classification, particles from the best-resolved class were re-extracted in a 480 px box and duplicate particles removed. Non-uniform refinement of these 71,430 particles was performed, before the detergent micelle and ECD were subtracted. To improve the density for MGRN1, the particles were locally refined with a mask encompassing MGRN1 and the TMD. This resulted in a map with an overall resolution of 4.41 Å. MEGF8-MOSMO-NB270 and MGRN1 local refinement maps were combined in PHENIX to create a composite map.

For the wild-type MEGF8^ΔR1R2ICD^-MOSMO-NB270 complex, 1.2 M particles were initially extracted in a 360 px box Fourier cropped to 120 px (2.191 Å/px) from 10,313 micrographs (**Figure S2D**). 2D templates were generated using 269,482 particles for template picking. Subsequently, one round of ab-initio coupled to heterogeneous refinement was performed on the unduplicated particles (628,472 images in the dataset). The best-resolved class containing 231,879 particles was used to train Topaz picking. 4 ab-initio models were generated with the picked particles and then refined using heterogeneous refinement. To increase the number of picked particles, an additional 1,844 movies were collected on the same grid. This dataset was processed in the same manner and extracted 77,574 particles in a 384 px box. The particles were then merged to create a refined set of 440,449 unbinned particles, before running a heterogeneous refinement. Those particles with a 3D class posterior probability lower than 90% in the best-resolved class (71,691 particles) were discarded. Non-uniform refinement was performed on the remaining particles (228,513 particles), resulting in a 3.03 Å map. The detergent micelle was masked out in a local refinement, which improved the map resolution to 2.98 Å.

### Note 2: Structural analysis of MOSMO

The ECD of MOSMO contains a β-sheet composed of five anti-parallel β strands, with β1-4 strands forming the extracellular segment 1 (ECS1) and β5 in ECS2 (**Figure 2A** and **Figure S3A**). The claudin W_28_-GLW_43_-C_46_-C_56_ consensus motif is located in the ECS1, with W43 being substituted by V43 in MOSMO ^98,99^. Trp28 is buried in a hydrophobic pocket formed at the top of the four-helix bundle, anchoring the ECS1 to the TMD. ECS1 is further stabilized by a conserved disulfide bond between Cys46 and Cys56, and a network of water molecules forming hydrogen bonds with residues in the extracellular loops connecting β1–β2 and β3–β4 strands (**Figures 2A-2C**). Furthermore, the Arg54 side chain forms three hydrogen bonds with residues in β2 and β3 strands of ECS1, limiting conformational flexibility of the ECD (**Figure 2B**). The shorter ECS2 consists of a loop region that connects to the β5 strand (**Figure 2C**). The analogous ECS2 loop region of claudins is less structured and more variable compared to MOSMO (**Figure S3B**). Gln129–Lys132 residues form a type II β-turn in MOSMO, stabilized by hydrogen bonds between the two residues (**Figure 2C**). In contrast, this same region is mainly unstructured in other claudins (e.g. Claudins-3, −4 and −19) (**Figure S3B**).

Both MOSMO and claudins possess an extended TM3 that protrudes into the extracellular space^56,57,98,100–102^ (**Figure S3B**). A proline residue located at the TM3 thenar position^56^ (e.g. Pro134 in mouse Claudin-3), conserved in approximately 50% of claudin family members, causes the top part of TM3, which protrudes outside the membrane, to bend away from the extracellular domain in claudins. However, in MOSMO, this conserved proline (Pro118) induces TM3 bending ∼20° toward the ECS1. This kink is stabilized by a backbone hydrogen bond between the carbonyl of Pro118 and backbone nitrogen of Phe121 in addition to hydrogen bonds between residues in TM2 and ECS2. This distinct orientation of the MOSMO TM3 is likely facilitated by its binding to MEGF8.

### Note 3: Structural analyses of MEGF8 Stem domain and its interface with MOSMO

The MEGF8 Stem domain, comprising 11 β-strands organized into three sheets, adopts a novel β-sandwich-like fold that groups with CUB modules by FoldSeek^103^ and Dali analysis^104^ (**Figures S3C to S3F**). Four anti-parallel (β1, β11, β3, and β8) strands and one parallel (β6) strand form one β-sheet. The second β-sheet, which forms one half of the Stem-MOSMO interface, is composed of three anti-parallel strands (β11, β2, and β9) and one parallel strand (β7). The two β-sheets pack against each other (**Figures S3D and S3E**) through hydrophobic interactions and hydrogen bonds, including the hydrogen bond between the Arg2500 side chain (β6) and Thr2512 carbonyl (β7). In addition, residues Tyr2533 (β9), His2538 and Phe2545 (β10) form π–π stacking interactions. An anti-parallel b-hairpin, composed of β4 and β5 strands, inserts into first β-sheet between residues Phe2444 and Ile2459, and is followed by a disordered proline-rich loop (residues Pro2463 to Glu2496) (**Figures S3C and S3E**). One extracellular domain (ECDII) of SIDT1 (PDB ID: 8JUL) shares a similar core fold to the MEGF8 Stem domain^58^ (**Figure S3E**). A comparison with the homologous, but more economical Stem domain from Attractin suggests a conserved 9 β-strand core closely related to prototypical CUB domains, like the *Clostridium histolyticum* collagenase CUB domain (PDB: 1NQJ)^105^ (**Figure S3F**). MEGF8’s Stem domain has a ∼53 amino acid Gly-Pro-Ser rich insert, which is unresolved in our cryo-EM structure, as well as an additional β-sheet (composed of β4 and β5 strands) that packs against the β-sheet that is not involved in MOSMO binding (**Figures S3D and S3F**). The MEGF8 Stem insert loop packs against a β-sheet that is not involved in MOSMO binding. The MEGF8 Stem-MOSMO interface shares striking similarities to the interaction interface between the CUB domain of *Clostridium perfringens* enterotoxin and Claudin-3^56^ (PDB: 6AKE, **Figure S3G**).

### Note 4: Structural analysis of MGRN1 engager domain

The MGRN1 engager domain adopts a β-sandwich fold (**Figure S3H**). FoldSeek^103^ and Dali analysis^104^ reveals that the MGRN1 engager domain clusters with ASF1 histone chaperone structures that have a core 7 β-strand fold related to Ig domains, with packed ABE and GFCD β-sheet faces^60^ (**Figures S3H and S3I**). MGRN1 has a 19 amino acid insertion between β-strands C and D, that is folded as a β-hairpin packed against its GFCD face.

## Supplementary videos

**Video S1: Structural overview of the MMM complex.**

The composite cryo-EM map of the ER/K-stabilized MMM complex (termed HelixMut) is shown within a low-pass filtered density map of the surrounding detergent micelle. A morph between the HelixMut and wild-type models reveals distinct orientations of the MEGF8 SAH. Close-up views highlight palmitoylation and sterol-binding sites, as well as the MEGF8 Stem-MOSMO and MEGF8 ICD-MGRN1 interfaces. Our structure-guided mutagenesis analysis (mutated residues marked by asterisks) validates structure features observed in the cryo-EM map and identifies key regions important for MMM function. Finally, superimposing the RNF4-E2-ubiquitin complex and MGRN1 RING domain better shows that E2-ubiquitin can be accommodated in the MMM complex without steric hindrance, supporting a plausible model for ubiquitin transfer at the membrane.

**Video S2: Molecular dynamics simulations of the MEGF8 SAH.**

All-atom molecular dynamic simulations of our “wild-type” and ER/K stabilized (“HelixMut”) cryo-EM structures of the MMM complex reveal the local motion of the MEGF8 TM-SAH. Despite both helices having an indiscernible degree of flexibility, only the active “wild-type” SAH moves towards the inner leaflet of the plasma membrane over time.

**Video S3: Molecular dynamics simulations of the MEGF8-SMO transmembrane interface.**

All-atom molecular dynamics simulations confirm that the MEGF8-SMO TM interface remains stable in POPC bilayer, supporting the AlphaFold-based MMM-SMO model prediction.

